# U6 snRNA m6A modification is required for accurate and efficient cis- and trans-splicing of *C. elegans* mRNAs

**DOI:** 10.1101/2023.09.16.558044

**Authors:** Aykut Shen, Katarzyna Hencel, Matthew T. Parker, Robyn Scott, Roberta Skukan, Aduragbemi S. Adesina, Carey L. Metheringham, Eric A. Miska, Yunsun Nam, Wilfried Haerty, Gordon G. Simpson, Alper Akay

## Abstract

pre-mRNA splicing is a critical feature of eukaryotic gene expression. Many eukaryotes use cis-splicing to remove intronic sequences from pre-mRNAs. In addition to cis-splicing, many organisms use trans-splicing to replace the 5′ ends of mRNAs with a non-coding spliced-leader RNA. Both cis- and trans-splicing rely on accurately recognising splice site sequences by spliceosomal U snRNAs and associated proteins. Spliceosomal snRNAs carry multiple RNA modifications with the potential to affect different stages of pre-mRNA splicing. Here, we show that m6A modification of U6 snRNA A43 by the RNA methyltransferase METT-10 is required for accurate and efficient cis- and trans-splicing of *C. elegans* pre-mRNAs. The absence of U6 snRNA m6A modification primarily leads to alternative splicing at 5′ splice sites. Furthermore, weaker 5′ splice site recognition by the unmodified U6 snRNA A43 affects splicing at 3′ splice sites. U6 snRNA m6A43 and the splicing factor SNRNP27K function to recognise an overlapping set of 5′ splice sites with an adenosine at +4 position. Finally, we show that U6 snRNA m6A43 is required for efficient SL trans-splicing at weak 3′ trans-splice sites. We conclude that the U6 snRNA m6A modification is important for accurate and efficient cis- and trans-splicing in *C. elegans*.

## Introduction

Recognition of 5′ and 3′ splice sites (SS) and the branchpoint sequence is crucial for the efficiency and accuracy of pre-mRNA splicing. How the spliceosome recognises the correct splice sites remains an important question in biology. In humans, 5′SSs are generally identified first by U1 snRNA through base pairing (1, 2), which is followed by the U5 snRNA binding to the 5′SS exon sequences −3 to −1 (3, 4), and U6 snRNA binding to the 5′SS intron sequences +3 to +5 (5). The 3′SS recognition is facilitated by the coordinated binding of the U2AF65 - U2AF35 heterodimer to the 3′SS and the SF1/BBP and U2 snRNA to the branch point sequence (6, 7). Following the recognition of 5′ and 3′SSs, U6 snRNA forms a helix with the U2 snRNA, leading to the formation of the B complex (8–11).

Splice site recognition is generally well-conserved among organisms with some differences. In *Caenorhabditis elegans (C. elegans)*, only a small number of 5′SSs have the AG//GUAAG sequence that would form a continuous Watson-Crick base pairing with the U1 snRNA as well as interacting strongly with the U5 and U6 snRNAs (12, 13). Furthermore, *C. elegans* and likely other nematodes do not have a branchpoint consensus (14). Interestingly, *C. elegans* SF1/BBP retained the conserved branch point recognition domain, and the *C. elegans* U2 snRNA has the same conserved antisense sequence found in mammalian U2 snRNA that matches the branch point motif (15). Therefore, it is unclear how SF1/BBP and U2 snRNA bind to branch point sequences in *C. elegans. C. elegans* introns have a highly conserved UUUCAG/R 3′SS sequence (Figure S1A). The short UUU sequence likely acts as the polypyrimidine tract recognised by the U2AF65, and U2AF35 recognises the CAG/R sequence for efficient splicing (16).

How U5 and U6 snRNA binding preferences affect 5′SS selection remains unclear. In *S. cerevisiae,* the U5 and the U6 snRNAs base pair with the largely invariant 5′SS sequences and stabilise the B-complex formation (10). However, metazoan 5′SSs have degenerate sequences. Although U5 and U6 snRNA interactions at the 5′SS are conserved (5), they form less continuous Watson-Crick base pairing, suggesting other factors could play a role in stabilising U5 and U6 snRNA interactions with the pre-mRNA. Several modifications found on snRNAs could contribute to snRNA - splice site interactions while recognising variable splice site sequences (17). For instance, the U5 loop I sequence is modified to UUmUψ (Um- 2′O- methyl, ψ-pseudouridine) (18), but the role of these modifications is not clear. U6 snRNA is m6A modified at A43 in many organisms by the conserved RNA methyltransferase METTL16 (human) / FIO1 (*A. thaliana)* / METT-10 *C. elegans)* / MTL16 (*S. pombe)* (19–24). Mutations in the *S. pombe* MTL16 and *A. thaliana* FIO1 cause numerous splicing errors (21, 24). In addition, the recently characterised U6 snRNA m2G72 modification also plays an important role during pre-mRNA splicing in human cells (25).

In addition to cis-splicing, nematodes, flatworms, cnidarians, rotifers, euglenozoa, and urochordates use spliced leader (SL) trans-splicing for mRNA maturation (26, 27). SL trans-splicing replaces the 5′ ends of pre-mRNAs with the non-coding SL RNA (28). SL trans-splicing uses the same snRNAs U2, U4, U5 and U6, as in cis-splicing, except for U1 (29, 30). In SL trans-splicing reactions, 5′ and 3′SSs are split between separate RNA molecules; the 5′SS resides within the SL RNA, and the 3′SS is generally positioned within the 5′ untranslated region of the pre-mRNA. Therefore, the trans-spliceosome needs to bring two separate RNA molecules together. The majority of the trans-spliced RNAs use SL1 non-coding RNA (80-90% of all trans-spliced RNAs), and the SL2 non-coding RNA is mainly used by transcripts in operons (7% of all trans-spliced RNAs) (31–33). The 5′SS on the SL1 RNA is an invariant AG//GUAAA, and most of the *C. elegans* 3′ trans-splice sites have the same UUUCAG/R sequence found in cis-spliced 3′SSs. The similarity of 5′ and 3′SSs between cis- and trans-splicing poses another challenge for the spliceosome during correct splice site recognition in *C. elegans*.

In *C. elegans*, the U6 snRNA m6A methyltransferase METT-10 / METTL16 is required for germ cell proliferation and larval development (34). Without *mett-10* function, germ cells arrest during mitosis and fail to enter meiosis, and animals develop multiple larval developmental defects, including embryonic lethality. Recent studies have implicated METT-10 in mRNA m6A modification and S-adenosylmethionine (SAM) homeostasis in *C. elegans* but failed to identify global pre-mRNA splicing defects (23, 35).

In order to understand the transcriptome-wide functions of METT-10 in *C. elegans*, we used short- and long-read sequencing to comprehensively analyse the cis- and trans-splicing of pre-mRNAs alongside their expression. Our results show that METT-10 loss-of-function causes global splicing defects in *C. elegans,* with both the cis- and the trans-spliceosomes failing to splice hundreds of genes accurately or efficiently. We identify the 5′SS +4A as a critical determinant of cis-splicing in the absence of U6 snRNA m6A43 modification in *C. elegans.* We further show that strong U5 and U6 interactions at the 5′SSs can support splicing at weak 3′SS sequences. As the SL RNA 5′SS is invariant, we show that most of the trans-splicing defects result from weak 3′SSs identified using U2AF binding motifs. Finally, we show that U6 snRNA m6A43 likely functions together with the spliceosomal protein SNRP-27 / SNRNP27K for accurate 5′SS recognition.

## Materials and Methods

### Nematode culture, strains, and maintenance

*C. elegans* strains were grown on Nematode Growth Medium (NGM) agar plates with *Escherichia coli (E. coli)* HB101 strain as a food source and maintained at 20°C unless stated otherwise. The following strains were used in the experiments: Wild type N2 Bristol, ALP010 *mett-10(ok2204*) III derived from backcrossing of VC1743 (34), ALP012 p*Mex-5*::*mett-10*::OLLAS::*tbb-2_UTR* (rna004) II generated using MosSCI (36). All experiments were performed starting from synchronous L1 animals, generated by hypochlorite treatment in a 2:2:1 solution (sodium hypochlorite (4-5%), H2O and 10M NaOH).

### Worm collection and RNA extraction

*C. elegans* strains N2 Bristol, ALP010, and ALP012 were grown on *E. coli* HB101 and collected at the young adult stage. RNA was isolated using TRIsure (Bioline, Cat. No. BIO-38032) following standard phenol-chloroform RNA extraction. The purity of the RNA was checked with Nanodrop, and the integrity was quantified using the Agilent 2200 TapeStation System. RNA concentration was measured with Qubit using RNA HS Assay (Invitrogen™ Q33224).

### Nanopore RNA Sequencing

100 µg of total RNA was used to perform poly(A)+ RNA isolation using PolyATtract® mRNA Isolation Systems (Promega UK LTD, Cat. No. Z5310). 750 ng of recovered mRNA was used for library preparation using the Direct RNA Sequencing kit following the manufacturer’s instructions (Oxford Nanopore Technologies, SQK-RNA002). Libraries were quantified using Qubit dsDNA HS Assay (Invitrogen™ Q32851). Sequencing was done on Oxford Nanopore Technologies MinION devices using flowcells R9.4.1.

### Illumina RNA Sequencing

1 µg of total RNA was used to perform mRNA isolation using NEBNext Poly(A) mRNA Magnetic Isolation Module (NEB Cat. No. E7490). The resulting mRNA material was used to prepare the libraries with the use of NEBNext® Ultra II Directional RNA Library Prep Kit for Illumina® (NEB, Cat No. E7760S) following the manufacturer’s instructions. Libraries were quantified using Qubit dsDNA HS Assay (Invitrogen™ Q32851). Sequencing was done at the Novogene (UK) Company Limited using an Illumina NovaSeq 6000.

### Egg-laying assay

Synchronous L1 animals were plated on *E. coli* HB101-seeded NGM plates at 20°C. Animals were grown to the larval stage L4 and shifted to 15°C, 20°C and 25°C. Each animal was transferred to a new plate twice daily. Egg-laying was performed with 1 technical replicate per biological replicate (genotype); 4 animals for N2 and ALP012 and 8 for ALP010 were used. Eggs were counted for 3 days using biological triplicates. One-way ANOVA was used to test significance in the egg-laying assay, followed by a Tukey’s post-hoc test. Normality and homogeneity of variances were, respectively, assessed by Shapiro-Wilk and Leven tests.

### Developmental assay

Synchronous L1 stage animals were individually picked onto *E. coli* HB101 seeded NGM plates and exposed to different temperatures. The developmental stages of animals were determined in biological triplicate using a stereomicroscope 42 hours post L1 and classified as L2-L3, late L4 and young adult.

### Nanopore Direct-RNA sequencing data processing

Read signals were base called using Guppy version 5.0.11 in GPU mode, employing the high-accuracy RNA model with the parameters -x cuda:all:100%, --num_callers 24, --calib_detect, --reverse_sequence yes, and -c rna_r9.4.1_70bps_hac.cfg. For mRNA modification analysis with Yanocomp (37), reads were aligned to the WBcel235 *Caernohabditis* reference transcriptome and the *de novo* assembled transcriptome using minimap2 version 2.17 (38) with the following parameters: -a -L –cs =short k14 –for-only –secondary =no. For other analyses, reads were aligned to the WBcel235 *Caernohabditis* reference genome using a two-pass alignment approach with minimap2 (38) and 2passtools version 0.3 (39). For transcriptome assembly and differential error site analysis, the first pass alignments were generated with the following parameters: --splice, -k 14, -uf, -w 5, -g 2000, -G 200000, --end-seed-pen 15, -A1, -B2, -O2,32, -E1,0, -C9, --splice-flank=yes, and -z200. High-confidence splice junctions from each replicate were extracted using 2passtools score and then merged using 2passtools merge to create a final set of trusted splice junctions. In the second pass, the reads were realigned to the WBcel235 reference genome using minimap2 with the same parameters as before, in addition to using the trusted junctions as a guide with the parameters -junc-bed and -juncbonus=10. For outron retention analysis, the first pass alignments were generated using minimap2 with parameters: -ax splice, -uf, -L, --cs, -k 14, and -G 50370. High-confidence splice junctions were extracted using the 2passtools score and merged into a final set of trusted splice junctions using 2passtools merge. Second-pass alignments were generated using the following parameters: -ax splice, -uf, -L, --cs, -k 10, and -G 50370, along with the trusted junctions from the first pass Nanopore direct RNA sequencing alignments.

### Nanopore poly(A)+ mRNA modification analysis

Two computational methods were used to identify METT-10 dependent RNA modifications: First, differential modification analysis was conducted at the signal level using the ‘n-sample’ GitHub branch of Yanocomp (37), by aligning the reads to the WBcel235 *C. elegans* reference transcriptome or the *de novo* assembled transcriptome, followed by generation of kmer-level signal data using f5c event align version 0.7 (40, 41) and Yanocomp prep. A three-way comparison of the genotypes was performed using the Yanocomp gmmtest, with a minimum KS statistic of 0.25. A false discovery rate (FDR) of 0.05 was applied as a cutoff to identify transcriptomic sites with significant changes in the modification rate. For the second method, differential error site analysis was conducted using “differ” (42), by aligning the reads to the genomic sequence using 2-pass alignment (as described above) and pairwise comparisons were conducted for different combinations of genotypes (wild type, ALP010, and ALP012) with a median expression threshold of 5 and with or without CPM normalisation. An FDR of 0.05 was applied as a cutoff to identify genomic sites with significant changes in the error rate.

### Identification of trans-spliced sites

Two different methods were employed to identify trans-spliced sites: 1) Detection from Nanopore direct RNA sequencing data through pairwise alignment of 5’ ends to the SL leader sequence, and 2) Detection from Illumina RNA-seq data through SL leader sequence trimming(43). For the first method, wild type, ALP010, and ALP012 direct RNA sequencing data were used to predict nearby acceptor sites by searching for an “AG” motif that could serve as a trans-splicing site within 10 nt of the aligned 5’ end of reads. The known sequence of splice leaders (SLs) and the 24 nt genomic sequence downstream of the predicted acceptor site of the exon were concatenated to create an expected reference sequence for the trans-spliced mRNA. Smith Waterman local alignment of this reference sequence against the 5′ end of sequence of the nanopore direct RNA sequencing read (24 nt downstream of the aligned 5′ end plus up to 28 nt of the 5′ softclipped portion of the read) was then performed with the parasail alignment function sw_trace_striped_32 (44). For each read, the trans-splicing position and splice leader class (SL1 or SL2) with the best pairwise alignment score were retained. The quality of the pairwise alignment across the trans-splice site was assessed using the junction alignment distance, which is defined as the minimum distance from the splice junction to the first alignment error (39). Trans-splice sites where the junction alignment distance of the highest scoring read was greater than 8 nt, were retained as high-scoring putative trans-splice sites. These trans-splice sites were used to create a position specific scoring matrix (PSSM) for the outron motif which was then used to score all possible trans-splicing sites – a threshold for the PSSM score was determined by splitting the scores of all possible sites at deciles and identifying the decile that maximised the Chi^2^ statistic of the thresholded junction alignment distances. Finally, a high-confidence set of trans-splicing sites were generated by retaining positions with both a maximum junction alignment distance of 8, and a position specific matrix score above the threshold. For the second method, SL-like sequences were trimmed from the 5’ end of the forward reads of Illumina data using cutadapt with a minimum overlap of 7 nt, a maximum error rate of 0.09, and minimum length (after trimming) of 15 nt (43). For each read, the trimmed SL sequence and class (SL1 or SL2) was recorded. The reads were then aligned to the WBcel235 reference genome using STAR as described below. Positions with more than 10 aligned reads with a trimmed SL sequence were identified as candidates. False positives were filtered out by identifying the candidate sites where the upstream genomic sequence was identical to the trimmed SL sequence. High confidence sites were used to build a PSSM for the outron sequence and select a PSSM threshold, as described above. Sites with a minimum of 2 supporting reads and an outron sequence scoring above the threshold were retained as high-confidence trans-splicing sites. The sites from the Illumina and Nanopore methods were combined to generate a list of non-redundant trans-splice sites.

### Detection of METT-10 dependent trans-splicing defects

METT-10-dependent trans-splicing defects were detected by developing scripts to assess changes in trans-splicing levels per annotated trans-splice site between wild type, ALP010, and ALP012 genotypes pairwise. The annotated trans-splice site positions were assigned to gene features in Ensembl release 95 reference annotation using pybedtools version 0.9.0 (45, 46). In cases where sites overlapped with multiple genes, they were assigned to the genes whose 5’ exon boundaries showed more significant agreement with the annotated trans-splice site positions. In cases of a tie, trans-splice sites were assigned to both genes being compared. Overlapping reads with each annotated trans-splice position were identified using pysam version 0.21.0 (47). Reads were categorised as “trans-spliced” or “retained outron” based on whether the 5’ position of the read end fell within the −14 to +10 window of the trans-splice site or earlier (< −14), respectively. Reads categorised as “retained outron” were conditioned to have identical splice junctions immediately downstream of the trans-splice site as those categorised as “trans-spliced”. The relative proportions of each category were counted. Counts from replicates of the same genotype were aggregated, and a 2×2 contingency table was created for each annotated trans-splice site, comparing the two genotypes. Significantly altered trans-splicing profiles were identified by performing a G-test using scipy (48). For sites with a p-value of < 0.05, G-tests for homogeneity between replicates of the same genotype were conducted. Calculated p-values were adjusted for multiple testing using the Benjamini-Hochberg false discovery rate (FDR) method. Retained outrons that were also cis-spliced at the annotated trans-splice positions and did not have an annotated SL site on the upstream exon with the 5’ splice site were classified as “cis-spliced retained outron” (CSRO). Retained outrons that exhibited a higher difference in the proportion of trans-spliced (PSI) reads at a downstream SL site on the same exon were classified as “Alternative 3’ trans-splice site“. The remaining trans-splice sites that showed a significant difference in PSI were classified as “retained outron” (RO). Sequence logos per alternative splicing event were generated using matplotlib version 3.7.1 (49) and matplotlib_logo (https://github.com/mparker2/matplotlib_logo). Contingency tables of splice site classes at U2AF65 and U2AF35 interacting positions were generated by analysing the difference in positions (−6 to −4) of the 3’ trans-splice sites from the consensus motif UUU and the difference in positions (−3 to +1) of the 3’ trans-splice sites from the consensus motif CAGR, respectively. Heatmaps of contingency tables were generated using seaborn version 0.12.2 (50). Gene tracks, utilising reads aligned to the WBCel235 reference genome, were generated using pyBigWig version 0.3.18 (51), pysam version 0.21.0 (47), and matplotlib version 3.7.1 (49)

### Illumina RNA sequencing data processing

#### Genome alignment

Standard adapter sequences and common contaminants were removed from the paired-end reads using BBduk from the BBtools package version 37.62 (https://sourceforge.net/projects/bbmap/) with the following parameters: -k 21, -ktrim r, -hdist 1, -mink 11, -trimq 15, -qtrim rl, -minlen 35, -tpe, and -tbo. The quality of the reads was assessed using FastQC version v0.12.1 (https://www.bioinformatics.babraham.ac.uk/projects/fastqc/) and MultiQC version 1.13 (52). The reads were aligned to the WBcel235 *C. elegans* reference genome using STAR version 2.7.10b (53) with the following parameters: --outFilterMultimapNmax 3, --alignSJoverhangMin 8, --alignSJDBoverhangMin 3, --outFilterMismatchNmax 4, --alignIntronMin 39, -- alignIntronMax 20000, --chimOutType Junctions, --chimSegmentMin 15, -- chimScoreJunctionNonGTAG 0, and --chimSegmentReadGapMax 20000. A splice junction database was generated from the Ensembl release 95 reference annotation with the parameters --genomeSAindexNbases 12 and sjdbOverhang 149.

#### Transcriptome assembly

Condition-specific transcriptome assemblies were generated using Stringtie version 2.1.7 (54) from the pooled Illumina and complementary nanopore Direct-RNA sequencing alignments using the following parameters: --mix, --rf, -c 1, -s 1, -g 0, and -M 10. A unified set of transcripts was created by merging all resulting condition-specific assemblies with the Ensembl release 95 reference annotation using the Stringtie merge tool with parameters: -g 0, -F 0, -T 0, -f 0.001, and -i. Open reading frames were annotated using Transuite version 0.2.2 with parameters: --cds 50 and --ptc 90, which were used in outron retention analysis to assign SL sites to genes they likely originate.

#### Transcript quantification and alternative splicing

Transcript quantification and splicing analysis was done as previously described in Parker et al. 2022 (21). Assembled transcripts were quantified per Illumina RNA-seq sample using Salmon version 1.10.1 (55) with the parameters -l A and --validateMappings. The WBcel235 *C. elegans* reference genome assembly was used as a decoy. Local splicing events in the assembled annotation file were classified using SUPPA version 2.3 (56) with the parameters generateEvents, -f ioe, --pool-genes, and -e SE SS MX RI FL. Event-level relative abundance (PSI) values per sample for each local event were then estimated from the transcript-level quantifications with the parameters psiPerEvent and --total-filter 1. PSI values combined from all samples were loaded into Python version 3.8.12 using pandas version 1.0.1 (57–59). The relationship between genotype and PSI was tested by fitting generalised linear models (GLMs) per local splicing event using statsmodels version 0.11 (60). Calculated p-values were adjusted for multiple testing using the Benjamini-Hochberg false discovery rate (FDR) method. Local splicing events with significant changes in PSI between ALP010 and wild-type strains were identified using an FDR threshold of 0.05. Sequence logos per alternative splicing event were generated using matplotlib version 3.7.1 (49) and matplotlib_logo. Contingency tables of splice site classes at U5 and U6 interacting positions were generated using the difference of the −2 to −1 positions of the 5’SS from the consensus motif AG and the difference of the +3 to +5 positions of the 5’SS from the consensus motif RAG, respectively. Heatmaps of contingency tables were generated with seaborn version 0.12.2 (50). Gene tracks using reads aligned to the WBCel235 reference genome were generated using pyBigWig version 0.3.18 (51), pysam version 0.21.0 (https://github.com/pysam-developers/pysam) and matplotlib version 3.7.1 (49).

The effect size for each position around the 5′SS was calculated by taking the difference between the distribution of ΔPSI for splice sites which had a particular base at the position and the distribution of ΔPSI for splice sites which did not have the base. The significance was tested with a Wilcoxon signed-rank test. Where the resulting p-value was less than 0.01 the presence of a base at a particular site was determined to have a significant effect on the likelihood of an alternate splicing event occurring at the 5′SS splice site. The direction of the effect was calculated by comparing the proportion of splice sites with a given base at a given position between *mett-10^-/-^*and wild type to the proportion of splice sites without the given base that had a significant difference in alternate splicing. Where the proportion of events was higher in sites with the base, sites with that base at a given position were less favoured in the mutant and were given a negative sign. The signed significance of each base was plotted against the position in the 5′SS, with points sized according to the frequency at which the base occurred at a given position across all 5′SS in the genome.

#### Sequencing data

All RNA sequencing raw data has been deposited to the European Nucleotide Archive with the accession number PRJEB65287. RNA-Seq data used for the SNRP-27 is described in (61).

#### Recombinant METT-10 protein purification

Full-length *C. elegans* METT-10 was subcloned into the pET-21a vector for expression in *E. coli*. The construct was overexpressed in Rosetta (DE3) cells (Novagen) using autoinduction media (62). Soluble METT-10 was purified from soluble lysate using Nickel affinity chromatography (Ni-NTA) and further purified by ion-exchange chromatography and gel filtration chromatography.

#### In vitro methylation assay

The in vitro methylation assay was carried out as described by Wang et al. 2016 (63). Briefly, a 15 μL reaction mixture containing 50 mM Tris pH 8.5, 0.01% Triton X, 50 μM ZnCl2, 1 mM DTT, 0.2 U/μL RNasin, 1% glycerol, 1 μM [3H]-SAM (Perkin Elmer), 1 μM full-length *C. elegans* U6, and 750nM METT-10 was incubated at room temperature for 1 hr. The reaction mixture was blotted on Biodyne B nylon membranes and washed with buffer (20 mM Tris pH 7.5, 0.01% Triton X), deionised water, and 95% ethanol, in that order, and then subjected to liquid-scintillation counting using the TriCarb 2010 TR Scintillation Counter (Perkin Elmer). RNA levels with the incorporated 3H-methyl group are shown as disintegrations per minute (DPM). All in vitro methylation data are shown as mean ± SD from three replicates. The RNA substrate, *C. elegans* U6 RNA, was transcribed in vitro and purified after separating on a denaturing polyacrylamide gel (64). The template sequence encoding the *C. elegans* U6 is GTTCTTCCGAGAACATATACTAAA ATTGGAACAATACAGAGAAGATTAGCATGGCCCCTGCGCAAGGATGACACGCAAATTC GTGAAGCGTTCCAAATTTTT.

## Results

### Loss of METT-10 function leads to cis- and trans-splicing defects

*mett-10(ok2204) null* mutants (*mett-10^-/-^* hereafter) have a deletion removing the entire methyltransferase domain (34) and have been shown previously to lack any detectable U6 snRNA m6A43 modification (23). We further confirmed that a recombinant METT-10 protein can efficiently methylate *in vitro* transcribed U6 snRNA (Figure S1B). Next, we used Nanopore Direct RNA sequencing and Illumina paired-end short-read RNA sequencing to analyse isoform-specific gene expression and alternative splicing in *mett-10^-/-^* and wild-type animals (Figure 1A and 1B). Short-read sequencing generates high-coverage data for accurate quantification of cis-splicing. The Nanopore Direct RNA sequencing generates long-read data extending from the polyA-tail to the 5′ ends of transcripts, allowing the detection of individual RNAs that are either trans-spliced or not. We used three biological replicates per genotype of poly(A)+ RNA for the Nanopore Direct RNA Sequencing and four biological replicates per genotype for the Illumina sequencing. We obtained, on average, 2.8 million reads per replicate in Nanopore Direct RNA Sequencing with an average mapping efficiency of 99% and 50 million reads per replicate in Illumina sequencing with an average mapping efficiency of 98% (Table S1). We quantified the cis-splicing defects using Illumina reads (Figures 1A and C) and the trans-splicing defects using Nanopore reads (Figures 1B, and D). We mapped reads to a custom *C. elegans* reference transcriptome built using wild-type and *mett-10^-/-^* Illumina and Nanopore reads to capture novel transcript isoforms and splicing events. Comparison of cis-spliced RNA fractions for each transcript (Δpsi) between *mett-10^-/-^* and the wild-type animals identified, in total, 2456 differential cis-splicing events with a p-value < 0.05 or 1644 splicing events with an FDR < 0.05 (Figures 1C and 1E, Table S2).

**Figure 1.**
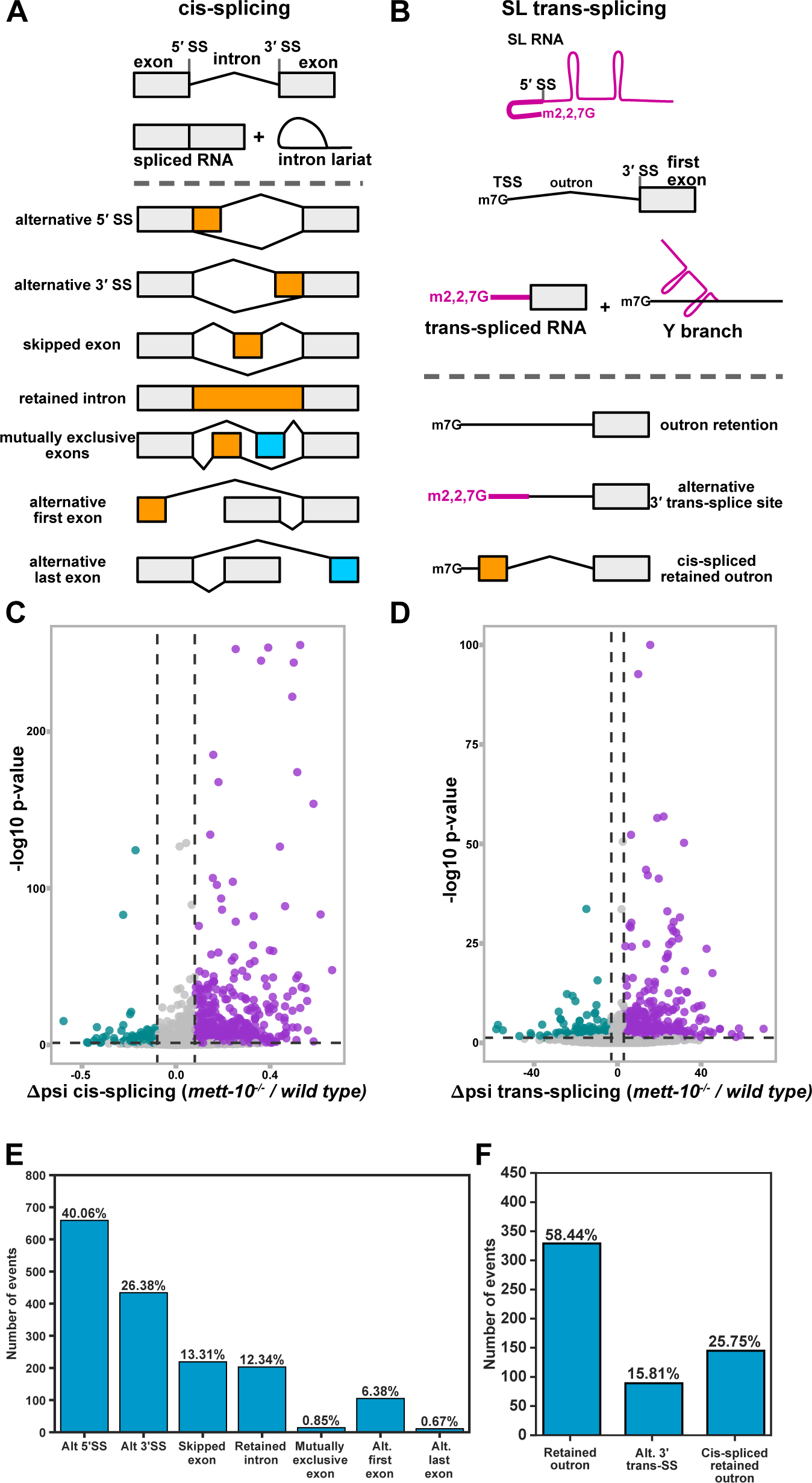
The absence of *mett-10* causes cis- and trans-splicing defects. **(A)** Overview of pre-mRNA cis-splicing and cis-splicing defects detectable by RNA sequencing. **(B)** Overview of pre-mRNA trans-splicing and trans-splicing defects detectable by RNA sequencing. **(C)** Volcano plot of all transcripts tested for cis-splicing changes between *mett-10^-/-^* and *wild-type* animals. Transcripts that show increased splicing in *mett-10^-/-^* (p ≤ 0.05 and ΔPSI ≤ −0.1) at a given splice site are coloured green, and transcripts that show reduced splicing in *mett-10^-/-^* (p ≤ 0.05 and ΔPSI ≥ 0.1) at a given splice site are coloured purple. **(D)** Volcano plot of all transcripts tested for trans-splicing changes between *mett-10^-/-^* and *wild-type* animals. Transcripts that show increased splicing in *mett-10^-/-^* (p ≤ 0.05 and ΔPSI ≤ −3) at a given splice site are coloured green, and transcripts that show reduced splicing in *mett-10^-/-^* (p ≤ 0.05 and ΔPSI ≥ 3) at a given splice site are coloured purple. -log10 p-value of 1 gene has been lowered to fit into the graph. **(E)** Classification of all significant (FDR < 0.05) cis-splicing defects in *mett-10^-/-^* animals**. (F)** Classification of all significant (p-value < 0.05) trans-splicing defects in *mett-10^-/-^* animals.

Out of 1644 cis-splicing events with an FDR < 0.05, the primary RNA splicing change in *mett-10^-/-^* animals is the alternative 5′SS usage (40.06%, Figure 1E). This is followed by alternative 3′SS usage (26.38%), exon skipping (13.31%), intron retention (12.34%) and alternative first exon usage (6.38%). In *A. thaliana,* the absence of U6 snRNA m6A43 leads to alternative 5′SS usage and intron retention events at similar levels, followed by alternative 3′SS choice and exon skipping (21). Therefore, the absence of U6 snRNA m6A modification in *C. elegans* leads to a stronger response in alternative 5′SS and 3′SS usage than other splicing events.

We next asked if trans-splicing was also affected in *mett-10^-/-^* animals. We quantified the trans-splicing defects using a new analysis pipeline using long nanopore direct RNA reads. First, we annotated the SL splicing sites using short and long-read data. Next, we counted reads containing the SL1 RNA or the outron sequence at the 5′-end of reads. Using this approach, we detected significant outron retention in 563 transcripts with a p-value < 0.05 or 232 transcripts with an FDR < 0.05 (Figure 1D and F, Table S3 and S4). Of 563 outron retention events with a p-value < 0.05, 58.44% are typical outron retained RNAs, and 25.75% are cis-spliced outron retained RNAs where the retained outron is cis-spliced, generating new exons. In addition, 15.81% of the outron retaining transcripts show alternative 3′ trans-splice site usage on the outron.

In summary, using a combination of short and long-read sequencing, we show that loss of METT-10 function causes a wide range of cis- and trans-splicing defects in transcripts from more than 2000 genes.

### 5′ splice-sites with +4A are sensitive to loss of METT-10 function

Sequence motif analysis of 5′SSs sensitive to loss of *mett-10* function shows that most of these splice sites have adenosine at the +4 position (+4A) within a //GURAG motif (Figure 2A, left panel). The alternative 5′SSs whose usage increases in *mett-10^-/-^* animals do not have +4A enrichment and are more likely to have an AG//GU splice site motif (Figure 2A, right panel, G-test p value = 1.19e-168). Next, we compared the frequency of 5′SS base composition for the bases that interact either with the U5 snRNA (Figure 2A, U5 class −1 and −2) or the bases that interact with the U6 snRNA (Figure 2A, U6 class +3, +4 and +5). The majority of the 5′SSs that are sensitive to the *mett-10* mutation have a non-AG U5 snRNA interacting sequence and an RAG for the U6 snRNA interacting sequence (Figure 2A, bottom left panel). In contrast, 5′SSs that are more often used in *mett-10^-/-^*animals are more likely to have an AG sequence for U5 snRNA interaction and a C, G or U at +4 position for U6 snRNA interaction (Figure 2A, bottom right panel). U5 snRNA loop I sequence interacts strongly with the exonic AG dinucleotides at the 5′SSs (3). Therefore, we see a shift from 5′SSs with a //GURAG motif and a weak U5 snRNA interacting sequence selected in wild-type animals to 5′SSs that lack the //GURAG motif but have a stronger U5 interacting sequence selected in *mett-10^-/-^* animals.

**Figure 2.**
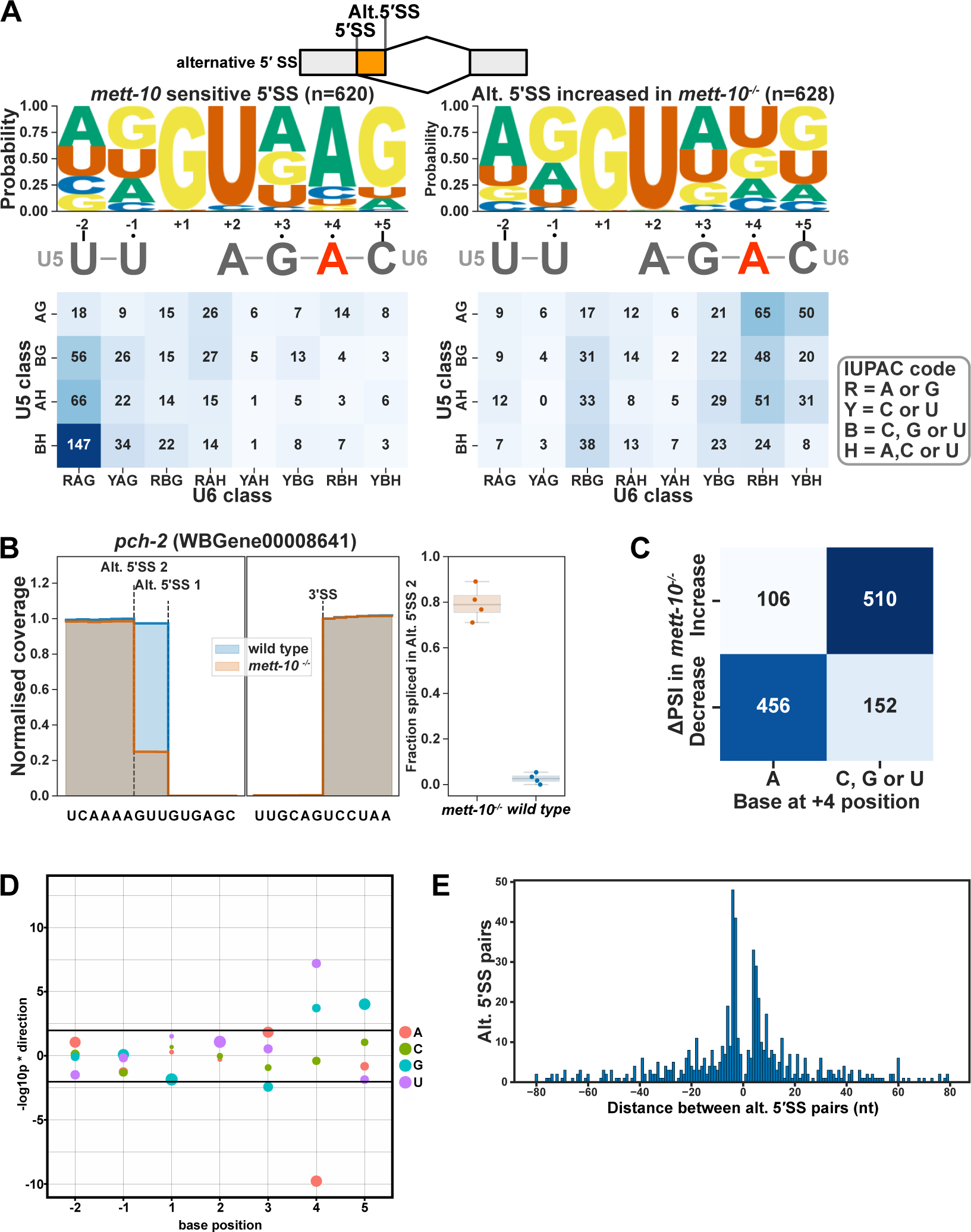
*mett-10* sensitive 5′ cis-splice sites have +4A. **(A)** The sequence motif and frequency analysis of *mett-10* sensitive 5′SSs (−2 to +5) and the alternative 5′SSs are used more often in *mett-10^-/-^*. The sequence motif shows the probability of bases at each position around the 5′SS. U5 and U6 snRNA binding sequences are shown under the sequence motif logo. The frequency table is coloured based on the U5 snRNA interacting sequence frequency on the y-axis (−2 and −1) and U6 snRNA interacting sequence frequency on the x-axis (+3,+4 and +5). **(B)** Normalised coverage of RNA-Seq reads for the *pch-2* intron 1 boundary. Alt. 5′SS 1 is used more often in wild-type animals and alt. 5′SS 2 is used more often in *mett-10^-/-^*. Barplot shows the fraction of reads supporting splicing at the Alt. 5′SS 2 over total reads in *mett-10^-/-^* and *wild-type* animals. **(C)** Heatmap showing the correlation between splice site usage in *mett-10^-/-^* (y-axis) and the specific base at position +4 of the 5′SS. **(D)** Effect size plot for the 5′SS positions −2 to +5. Negative values indicate bases at the specific position are associated with significantly more alternative splicing and positive values indicate bases at the specific position are associated with significantly less alternative splicing events. The size of the circles correspond to the frequency of the base at a given position across all 5′SSs in the genome. **(E)** Histogram for the distance between alternative splice site pairs. The Y-axis shows the number of alternative splice site pairs, and the x-axis shows the distance between the pairs, with negative values indicating the alternative splice site moves upstream and positive values indicating the alternative splice site moves downstream of the original splice site.

The global switches in 5′SSs can be detected at individual genes. For instance, at *pch-2* (pachytene checkpoint 2), which encodes the *C. elegans* orthologue of the human TRIP13 required for spindle checkpoint during mitosis and meiosis (65, 66), exon1 is exclusively spliced at the 5′SS UU//GUGAG in wild-type animals, whereas in *mett-10^-/-^*animals splicing occurs predominantly 3 nt upstream at the AA//GUUGU position (Figure 2B). As a result, PCH-2 amino acids Lys and Phe at positions 20 and 21 are replaced with an Asn. Similarly, at T12C9.7, which encodes the *C. elegans* orthologue of the human mitotic specific cyclin B2 (CCNB2), exon 8 is most frequently spliced at UU//GUGAG in wild-type animals and in *mett-10^-/-^* animals, 5′SS choice moves to the nearby UG//GUUG (Figure S2A, for additional examples see Figure S2A-C). Overall, 75% of all *mett-10* sensitive 5′SSs have +4A (Figure 2C). We calculated the effect size of each base, from −2 to +5 position, on alternative splicing of 5′SSs between *mett-10^-/-^* and wild-type animals by comparing the distribution of ΔPSI values of SSs with or without a given base at each position (Figure 2D). 5′SSs with +4A have a significantly higher percentage of alternative splicing, and 5′SSs with +4G, +4U or +5G are significantly less likely to be alternatively spliced (Figure 2D). In addition, in *mett-10^-/-^* mutants, alternative 5′SSs are equally found upstream or downstream and primarily within +/−5nt of the canonical splice site used in wild-type animals (Figure 2E). The alternative 5′SSs used in *A. thaliana* FIO1 mutants show a similar distribution (21), suggesting that the U1 snRNA selects multiple 5′SSs within a sequence window and the m6A modification status of U6 snRNA could determine the final 5′SS position.

In conclusion, METT-10 is required for the accurate splicing of 5′SSs with +4A, and the alternative 5′SSs used in the *mett-10^-/-^* mutants favour sequences without +4A and a stronger U5 interaction with the upstream exon sequences.

### Loss of METT-10 function leads to intron retention and exon skipping

We identified transcripts from 341 genes with p-value < 0.05 or 197 genes with an FDR < 0.05 that show altered intron retention levels in *mett-10^-/-^* animals compared to wild-type animals (Table S2). Out of 197 transcripts with an FDR < 0.05, 118 showed increased intron retention, and 79 showed reduced intron retention (Figure 3A). The 5′SSs of introns with increased retention in *mett-10* mutants have the //GURAG motif and weak U5 recognition sequence (Figure 3A, left panel). In contrast, the 5′SSs of introns with reduced retention have the AG//GU motif and lack +4A (Figure 3A, right panel, G-test p value = 2.9e-11). For example, at *C. elegans* gene Y18H1A.11, which encodes a choline-phosphate cytidylyltransferase, the mammalian orthologue of PCYT1, the 5′SS has CA//GUGAG sequence, which conforms to the //GURAG motif and a weak U5 interacting CA dinucleotide (Figure 3B).

**Figure 3.**
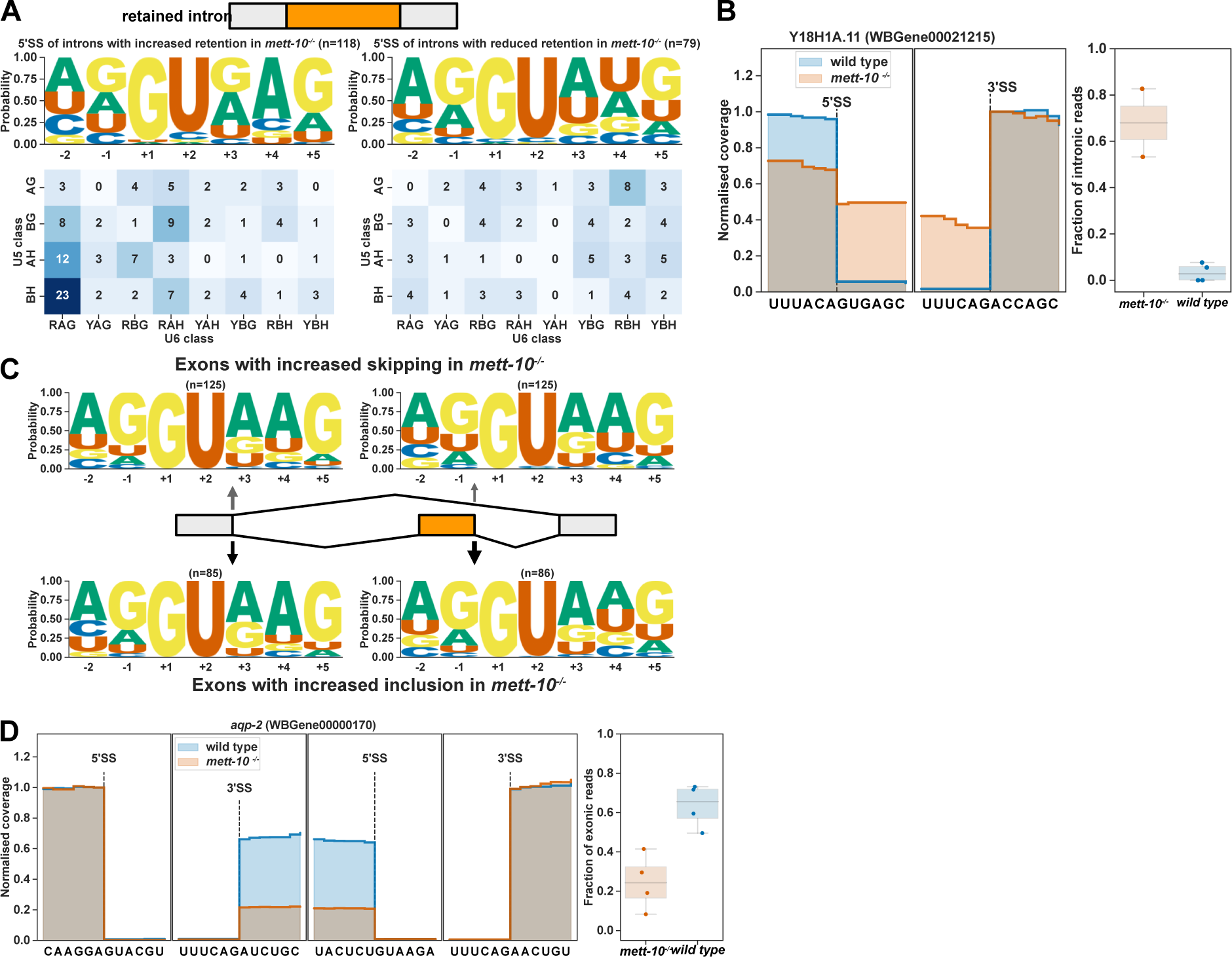
*mett-10^-/-^* animals have increased intron retention and exon-skipping. **(A)** 5′SS motif analysis of introns with increased (left) and decreased (right) retention in *mett-10^-/-^*. The frequency of sequences corresponding to U5 and U6 binding are shown in the heatmap. **(B)** Normalised RNA-Seq coverage of Y18H1A.11 intron 3 in *mett-10^-/-^*and *wild-type* animals. The bar plot shows the fraction of intronic reads over the retained intron. **(C)** 5′SS motif analysis of exons with increased skipping (upper panel) and increased retention (bottom panel) in *mett-10^-/-^*animals. Sequence motifs are shown for the upstream exon 5′SSs (grey) and the retained/skipped exon (orange). **(D)** Normalised RNA-Seq coverage of *aqp-2* exon 5 in *mett-10^-/-^* and *wild-type* animals. Bar plots show the fraction of exonic reads over the skipped exon.

Next, we compared the 5′SS sequences of introns adjacent to the exons that show either increased skipping or inclusion in *mett-10*^-/-^ animals compared to the wild-type (Figure 3C). Exons that show increased skipping have a 5′SS //GURAG motif with a higher probability of +4A compared to the exons that show reduced skipping, which have a higher probability of AG//GU (Figure 3C). However, the overall motif between −2 and +5 positions is not significantly different (G-test p-value=0.47). This could be due to multiple splice sites having an influence on the exon-skipping events. The exon 5 of *aqp-2*, the orthologue of the human Aquaporin3, is frequently skipped in *mett-10^-/-^* animals (Figure 3D). The 5′SS adjacent to the *aqp-2* exon 5 has the CU//GUAAG sequence that fits the //GURAG motif and a weak U5 interacting CU dinucleotide. Therefore, intron retention and exon skipping events observed without METT-10 function predominantly involve a +4A at 5′SSs.

### Loss of METT-10 function affects alternative 3′SS usage

The second most abundant class of cis-splicing changes observed in the absence of the METT-10 function is alternative 3′SS usage (Figure 1E). During splice site recognition, U6 snRNA interacts directly with the 5′SS sequence and, therefore, it is not expected to affect the 3′SS recognition by U2AF proteins or the branch-point recognition by the U2 snRNP. In *mett-10^-/-^* animals, we observed 656 alternative 3′SS usage events with a p-value <0.05, of which 434 had an FDR < 0.05 (Table S2). *mett-10* sensitive 3′SSs do not have the conserved *C. elegans* 3′SS motif of UUUCAG//R, which is required for efficient U2AF65-U2AF35 binding.

Whereas the 3′SSs with increased usage in *mett-10^-/-^* have a strong UUUCAG/R motif (Figure 4A, G-test −6 to +1 positions p-value=2.7e-149). In addition, most of the alternative 3′SS usage occurs at 5′SSs with a +4A (263), as opposed to non-+4A 5′SSs (167) (Figure S3A).

**Figure 4.**
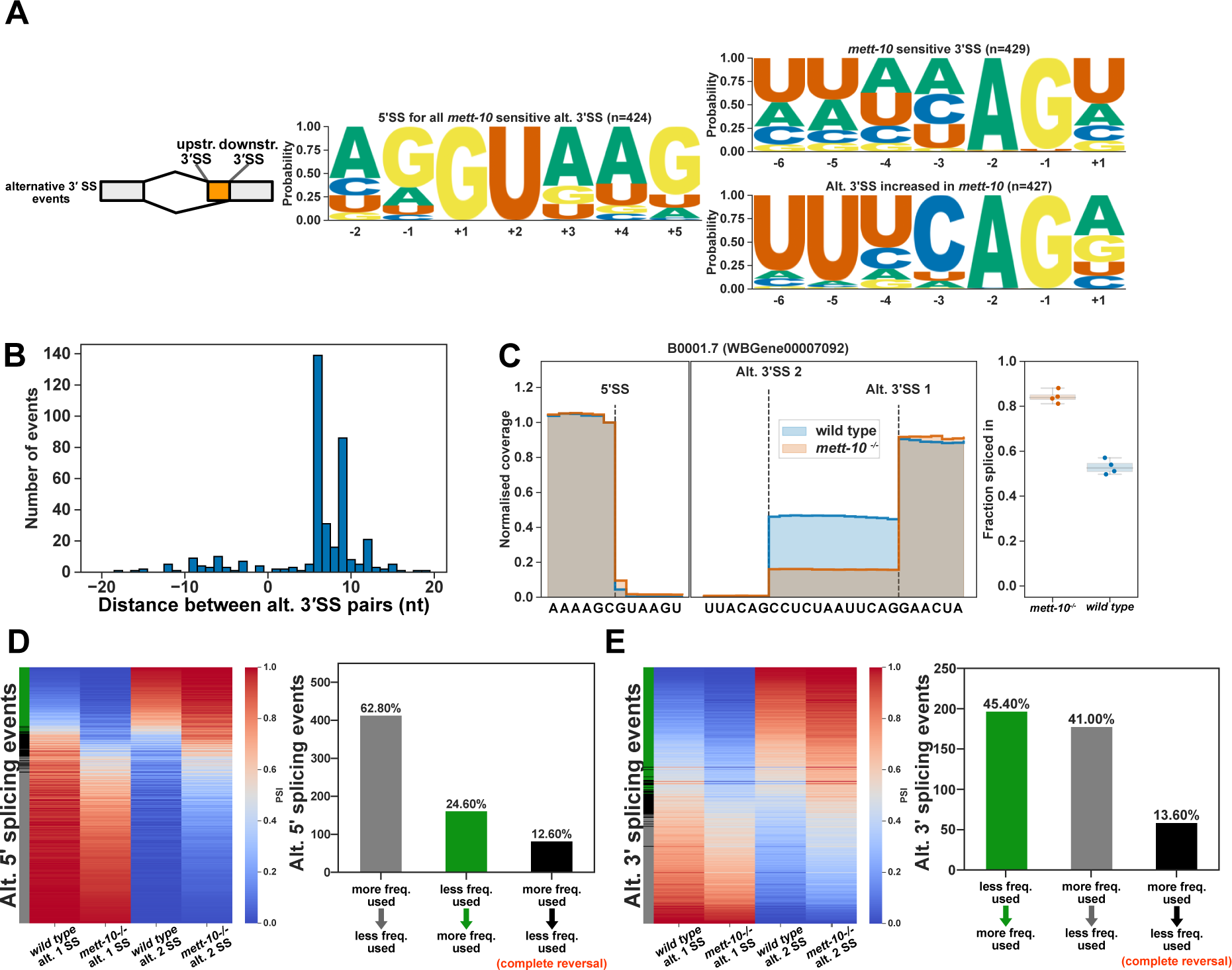
The absence of *mett-10* affects 3′SS usage. **(A)** 5′ and 3′SS motif analysis of transcripts that are *mett-10* sensitive (upper panel) and the corresponding alternative 3′SSs whose usage increases in *mett-10^-/-^* (bottom panel). **(B)** Histogram for the distance between alternative splice site pairs. The Y-axis shows the number of alternative splice site pairs, and the X-axis shows the distance between the pairs, with negative values indicating the alternative splice site moves upstream and positive values indicating the alternative splice site moves downstream of the original splice site. **(C)** Normalised RNA-Seq coverage of B0001.7 intron 6. The canonical splice position is Alt. 3′SS 1. In wild-type animals, weak upstream splice site Alt. 3′SS 2 is also utilised. The bar plot shows the fraction of reads covering the exon sequence to the right of the canonical splice site. **(D)** Heat map (left) and bar plots (right) showing the frequency of 5′SS usage at specific splice sites in wild-type and *mett-10^-/-^*animals. **(E)** Heat map (left) and bar plots (right) showing the frequency of 3′SS usage at specific splice sites in wild-type and *mett-10^-/-^*animals.

Unlike alternative 5′SSs, which can be found upstream and downstream of the canonical splice sites used in wild-type, alternative 3′SSs used in *mett-10^-/-^*animals are predominantly found downstream of the canonical 3′SS (Figure 4B). This contrasts with *Arabidopsis fio1* mutants, where alternative 3′SS usage can occur upstream or downstream from the canonical wild-type site (21). Most *mett-10^-/-^* sensitive 3′SSs (365) move from a weak upstream 3′SS sequence to a strong downstream 3′SS (Figure S3B, bottom panels). The remaining alternative 3′SS events (64) reflect shifts in usage from a stronger downstream 3′SS sequence to a moderate upstream 3′SS, where the +1R and −3C frequencies are lower than the 3′SSs shifting downstream (Figure S3B, top panels). For example, in wild-type animals, intron 6 of the B0001.7 gene is frequently spliced at the downstream 3′SS AUUCAG//G and the upstream 3′SS UUACAG//C (Figure 4C). In *mett-10* mutants, most of the splicing events occur at the downstream 3′SS and the frequency of splicing at the upstream 3′SS is significantly reduced (Figure 4C). Similarly, in wild-type animals intron 4 of *pdk-1*, a conserved protein kinase with a role in organismal sterility, is frequently spliced at the downstream 3′SS UUUCAG//A and the upstream 3′SS AGAAAG//U (Figure S2C). In *mett-10* mutants, the frequency of splicing events at the upstream 3′SS is significantly reduced, and most of the splicing happens at the downstream 3′SS (Figure S2C).

In contrast to shifts in alternative 5′SS usage in *mett-10* mutants from a site used more frequently in wild-type animals to a splice site used less frequently, alternative 3′SS usage shifts from a splice site used less frequently in wild-type animals to a splice site used more frequently in wild-type animals (Figure 4D and 4E). For the alternative 5′ and 3′SS usage observed in *mett-10* mutants, 12.6% of the 5′SS events and 13.6% of the 3′SS events show a complete reversal of the splice site usage from a more frequently used splice site in wild-type animals to a less frequently used splice site wild type animals (Figure 4D and 4E, black bars vs grey bars).

Mammalian METTL16 can methylate the pre-mRNA of S-adenosylmethionine (SAM) synthetase MAT2A at a UACAGA motif that is present within a hairpin structure (19, 20, 67–70). *C. elegans* METT-10 can also methylate the pre-mRNAs of three SAM synthetase genes, *sams−3, −4* and *−5,* at an intronic UACAG//A sequence that forms a hairpin (23, 35). METT-10 mediated m6A modification of *sams−3,−4 and −5* pre-mRNAs has been reported to cause the intron retention and alternative 3′SS usage of the *sams* genes in wild-type animals (23, 35). We observed similar intron retention and alternative 3′SS usage in *sams−3 and −4* except for *sams−5*, but in all cases, the amplitude of the intron retention and alternative 3′SS usage in wild-type animals were smaller than previously reported (Figure S4A and S4B). Of all the alternative 3′SS usage events that shift from an upstream position to the downstream canonical position (365), only 3.8% (14) have the methylation motif UACAG//A at the canonical splice site.

Oxford Nanopore Direct RNA sequencing data can predict RNA modifications (37, 71–74). Therefore, we used a signal-level analysis approach, Yanocomp (37), and a differential error rate approach, *differ* (42), to predict the modified bases in our Direct RNA Sequencing data. Our analysis of the Nanopore Direct RNA reads comparing the *mett-10* mutant and wild-type animals and using two different approaches did not reveal any significant events that can be attributed to RNA modifications at the *sams−3, −4* and *−5* intron sequences (Table S5 and S6).

### METT-10 is required for efficient SL trans-splicing

In addition to cis-splicing defects, we observed significant SL trans-splicing defects in *mett-10^-/-^* animals using Nanopore Direct RNA sequencing (Figure 1F). Although U6 snRNA was shown to be required for SL trans-splicing (29), nothing is known about the role of snRNA modifications in SL trans-splicing in general. To further examine the role of *mett-10* and U6 snRNA m6A43 in SL trans-splicing, we generated a single-copy insertion transgene expressing METT-10 protein under the control of a germline promoter in the *mett-10^-/-^*background (*mett-10 rescue).* As expected, germline expression of *mett-10* significantly rescues the fertility defects in *mett-10^-/-^* animals at all temperatures tested (Figure S5A). Surprisingly, germline expression of *mett-10* in *mett-10^-/-^* animals also rescues somatic developmental defects (Figure S5B), and animals appear like wild-type. Further supporting the phenotypic data, most trans-splicing events are rescued in *mett-10 rescue* animals (Figure 5A - D). Normalised coverage data of Nanopore Direct RNA sequencing reads shows that all trans-splicing classes, outron retention (Figure 5B), alternative 3′ trans-splice site usage (Figure 5C) and cis-spliced retained outron (Figure 5D) are effectively rescued. These results strongly support a role for METT-10 and U6 snRNA m6A43 in SL trans-splicing.

**Figure 5.**
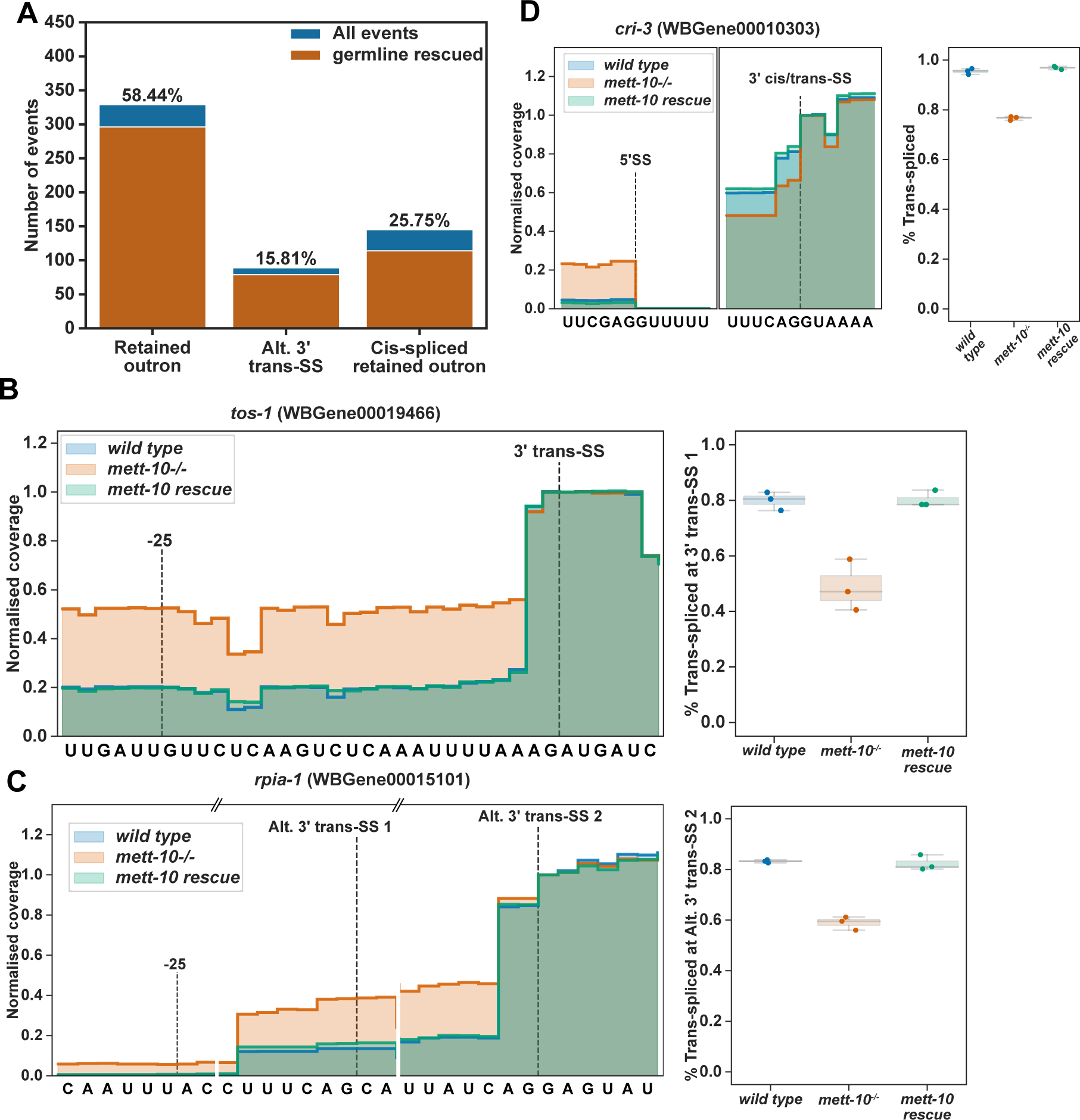
*mett-10* is required for efficient trans-splicing. **(A)** The trans-splicing defects rescued by the germline expression of *mett-10* are shown in orange over the total SL trans-splicing defects observed in *mett-10^-/-^* animals as in Figure 1F (blue). **(B - D)** Examples for the outron retention **(B)**, alternative 3′ trans-splice site usage **(C)** and cis-spliced outron retention **(D)** events showing the normalised RNA-Seq read coverage (left) and fraction of reads covering the outron sequence (right) in *mett-10^-/-^*and *wild-type* animals. Due to the sequence similarity between the SL1 RNA and the 3′ trans-splice sites, RNA-Seq read coverage drop does not always align with the actual splice site.

### 3′ trans-splice site sequence features determine *mett-10* sensitivity

U6 snRNA is essential for SL trans-splicing (29), but how it interacts with the 5′SSs on SL RNAs is not fully resolved. In *mett-10^-/-^*animals, although hundreds of genes have trans-splicing defects, most of the RNAs are still effectively trans-spliced. Therefore, we hypothesised that there could be sequence determinants of *mett-10^-/-^* trans-splicing sensitivity at the 3′ trans-splice site sequences rather than the invariable 5′ trans-splice sites. We grouped the transcripts with a significant trans-splicing change according to the extent of the trans-splicing difference (□PSI) and analysed the 3′ trans-splice site motifs separately (Figure 6A and S6). Compared to the 3′ trans-splice site motif of a background set of effectively trans-spliced genes, genes that show trans-splicing defects in *mett-10^-/-^* have weaker 3′ trans-splice site sequences (Figure 6A). Genes with lower □PSI have 3′ trans-splice site motifs with minor deviations from the background motif, particularly affecting the −4U and −3C and +1R (Figure 6A, middle panel and S6). Genes with higher □PSI show the most significant variation from the conserved UUUCAG/R motif, mainly at the −4U, −3C and +1R (Figure 6A, bottom panel and S6). The U2AF65-U2AF35 dimer recognises the 3′ trans-splice site sequence UUUCAG/R, with U2AF65 recognising the UUU sequence acting as a polypyrimidine track and the U2AF35 recognising the CAG/R sequence (16). We then analysed the frequency of U2AF65 and U2AF35 binding sequences at the 3′ trans-splice sites of the control sequences and the sequences that show significant □PSI (Figure 6B). Our analysis shows that both U2AF65 and U2AF35 recognition motifs deviate from the consensus UUUCAG/R motif, with the U2AF35 recognition sequence showing the largest deviation (Figure 6B).

**Figure 6.**
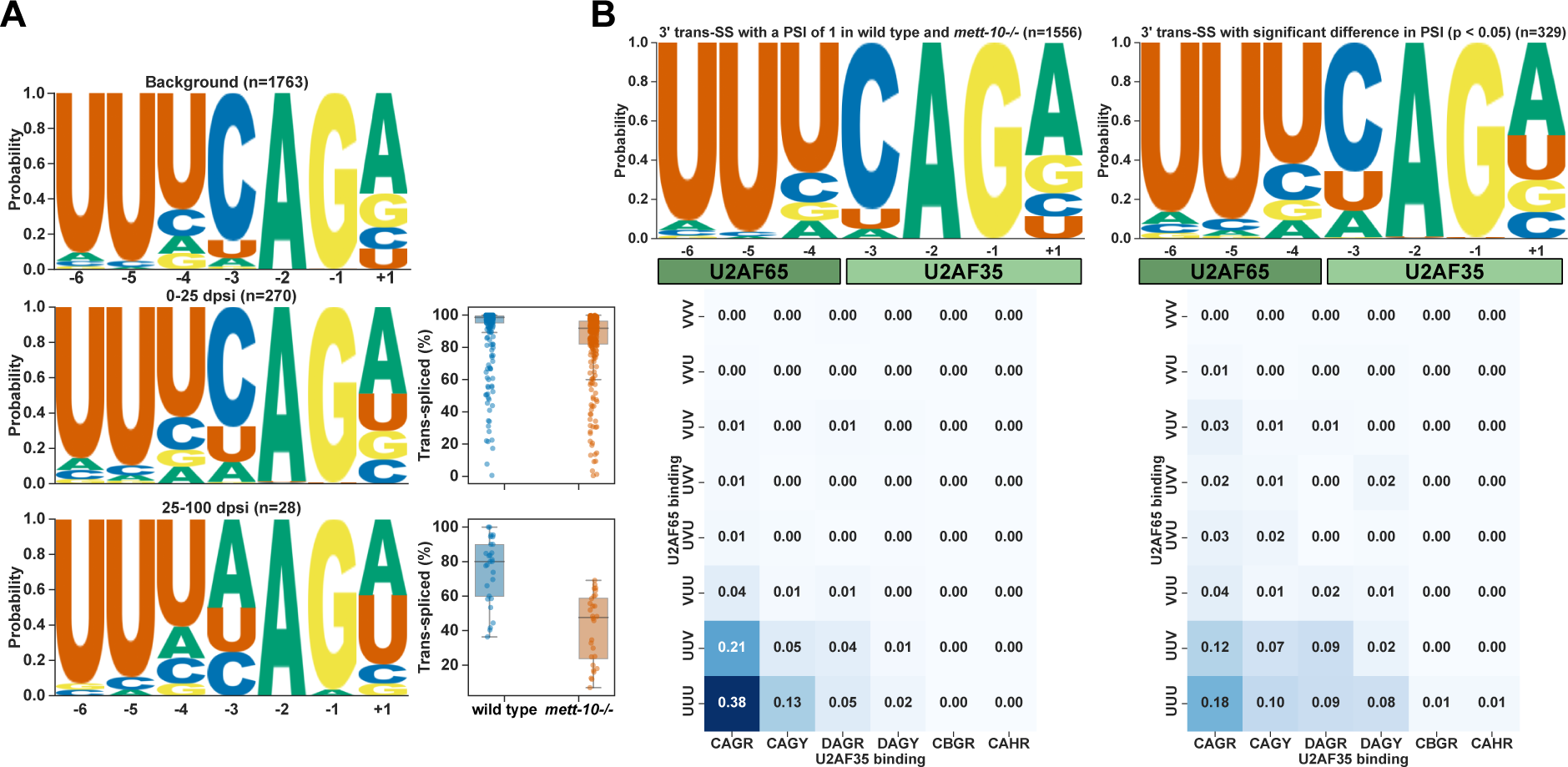
*mett-10* sensitive 3′ trans-splice sites have weak U2AF binding motifs. **(A)** 3′ trans-splice site motif of background transcripts that do not show the trans-splicing defect (top panel), transcripts that show the weak trans-splicing defect (middle panel) and transcripts that show the strong trans-splicing defect (bottom panel). **(B)** Frequency of sequences corresponding to U2AF65 binding (y-axis) and U2AF35 binding (x-axis) alongside the sequence logo of transcripts that do not show trans-splicing defect (left panel) and transcripts that show significant trans-splicing defect (right panel).

Our results show that without *mett-10*, *C. elegans* pre-mRNAs with weak 3′ trans-splice sites fail to trans-splice effectively and accurately.

### Interaction of U6 snRNA m6A43 and SNRNP27K in splice site selection

Spliceosomal snRNAs interact with many spliceosomal proteins during different stages of pre-mRNA splicing. Structural studies of spliceosomal complexes have not fully resolved how U6 snRNA m6A43 interacts with the 5′SS or the nearby proteins. A recent phylogenetic analysis of 5′SSs in *Saccharomycotina* identified a strong association between the conservation of METTL16 and the spliceosomal protein SNRNP27K (*snrp-27* in *C. elegans)* with the 5′SS +4A (75). Furthermore, M141T mutation in SNRP-27 leads to alternative splicing of 5′SSs with +4A (61). We re-analysed the previously published RNA-Seq data from *snrp-27(az26)* animals that carry the M141T mutation and the corresponding *wild-type* animals (61) using our RNA splicing analysis pipeline. We identified more alternative splicing events (2159) than previously reported, and the most abundant event class is alternative 5′SS usage (Figure 7A). Similar to *mett-10^-/-^*, *snrp-27(az26)* affects 5′SSs with +4A and a //GURAG motif, and the splicing happens at an alternative 5′SS with an AG//GU motif (Figure 7B). 60.7% of all 5′SS with a reduced usage in *snrp-27(az26)* have +4A (Figure S7A). The effect size comparison of each base, from −2 to +5 position, between *snrp-27(az26)* and wild-type animals shows that 5′SSs with +3G and +4A have a significantly higher percentage of alternative splicing in *snrp-27(az26)* and 5′SSs with -2A, +3A, +4U or +5G are significantly less likely to be alternatively spliced in *snrp-27(az26)* (Figure 7C). Similar to *mett-10* mutants, in *snrp-27(az26),* alternative 5′SSs can be found upstream and downstream of the canonical wild-type splice site (Figure S7B).

**Figure 7.**
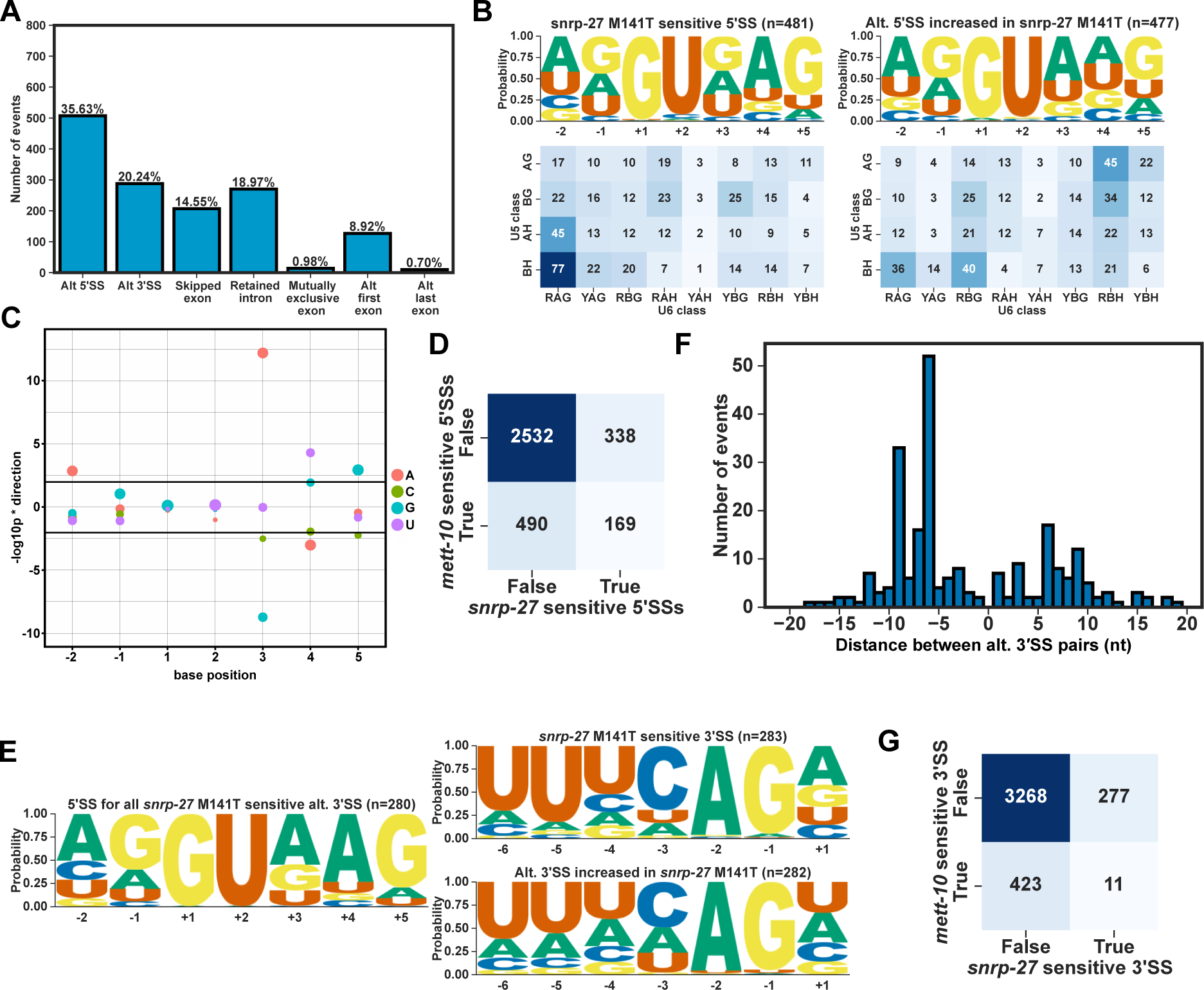
*snrp-27* is required for the recognition of 5′SSs with +4A. **(A)** Classification of all significant (FDR < 0.05) cis-splicing defects in *snrp-27(az26)* M141T animals compared to wild-type**. (B)** Sequence motif (upper panels) and the frequency of U5 and U6 interacting sequences (bottom panels) of the 5′SSs that are either sensitive to *snrp-27(az26)* (left panels) or used more often in *snrp-27(az26)* (right panels). **(C)** Effect size plot for the 5′SS positions −2 to +5. Negative values indicate bases at the specific position are associated with significantly more alternative splicing and positive values indicate bases at the specific position are associated with significantly less alternative splicing events. The size of the circles correspond to the frequency of the base at a given position across all 5′SSs in the genome. **(D)** Heat map showing the overlap of *mett-10* and *snrp-27* sensitive 5′SSs. **(E)** Sequence motif analysis of 3′SSs that are either sensitive to *snrp-27(az26)* (upper panel) or used more often in *snrp-27(az26)* (bottom panel). 5′SS motif of sensitive 3′SSs are shown on the left. **(F)** Histogram showing the distance between alternative 3′SS pairs (x-axis) and the number of 3′SS events (y-axis) **(G)** Heat-map showing the overlap of *mett-10* and *snrp-27* sensitive 3′SSs.

Overall, *mett-10* and *snrp-27* sensitive 5′SSs overlap substantially (Figure 7D). Although 5′SSs sensitive to either one or both proteins share a //GURAG motif, 5′SSs sensitive to both *mett-10* and *snrp-27* have a more dominant //GUGAG motif (Figure S7C). In contrast, 5′SSs that are only sensitive to *mett-10* have a //GUAAG motif (Figure S7D), and the 5′SSs that are only sensitive to *snrp-27* have a //GUDAG motif (D = A, G or U) (Figure S7E). Therefore, in addition to the +4 position, the +3 position of 5′SSs is also essential for effective and accurate splicing mediated by SNRP-27.

Unlike *mett-10* sensitive 3′SSs that are defined by a weak 3′SS motif, *snrp-27* sensitive 3′SSs have a strong UUUCAG/R motif, and the alternative 3′SSs appear to be weaker, despite having a similar //GURAG motif at their 5′SSs (Figure 7E). Furthermore, and in contrast to the alternative 3′SS usage in *mett-10^-/-^*, most of the alternative 3′SS events in *snrp-27(az26)* are found upstream of the canonical wild-type site (Figure 7F), and the shift happens from a strong UUUCAG/R sequence to a weak 3′SS motif (Figure S7F). These differences between *mett-10* and *snrp-27* sensitive 3′SSs are reflected in the lack of overlap in 3′SSs sensitive to the absence of either gene function (Figure 7G).

In summary, our results show that METT-10 and SNRP-27 are required for effective splicing of 5′SSs with +4A, and each protein affects a subset of genes with differences in their +3 nucleotide composition. In contrast, METT-10 and SNRP-27 affect distinct 3′SSs. Therefore, METT-10 and SNRP-27 likely play related but distinct roles in 5′SS selection.

### Consequences of cis- and trans-splicing defects in coding potential of transcripts

The alternative 5′ and 3′SS usage can add or remove in-frame amino acids or stop codons in the case of intron retention and exon skipping. The distribution of alternative 5′SSs in *mett-10^-/-^* lacks apparent 3nt periodicity seen in the distribution of alternative 3′SSs (Figure 2D and 4B). We analysed all cis-splicing events for their 3nt periodicity and categorised the events in-frame if the change is a multiple of 3nt and out-of-frame if the change is not a multiple of 3nt. Most intron retention events create out-of-frame changes, and most exon-skipping events create in-frame changes (Figure 8A). Most alternative 5′SS events are out of frame, whereas most 3′SS events are in-frame (Figure 8A). For instance, the alternative 5′SS in the neuronal transcript Y41C4A.12 creates an early termination codon truncating the open reading frame by 23 amino acids (Figure S8A). *C. elegans* C55A6.10 encodes a protein homologous to the human C12orf4, linked to autosomal recessive intellectual disability (76). The alternative 5′SS in transcripts of C55A6.10 creates an early termination codon in the middle of the gene (Figure S8B).

**Figure 8.**
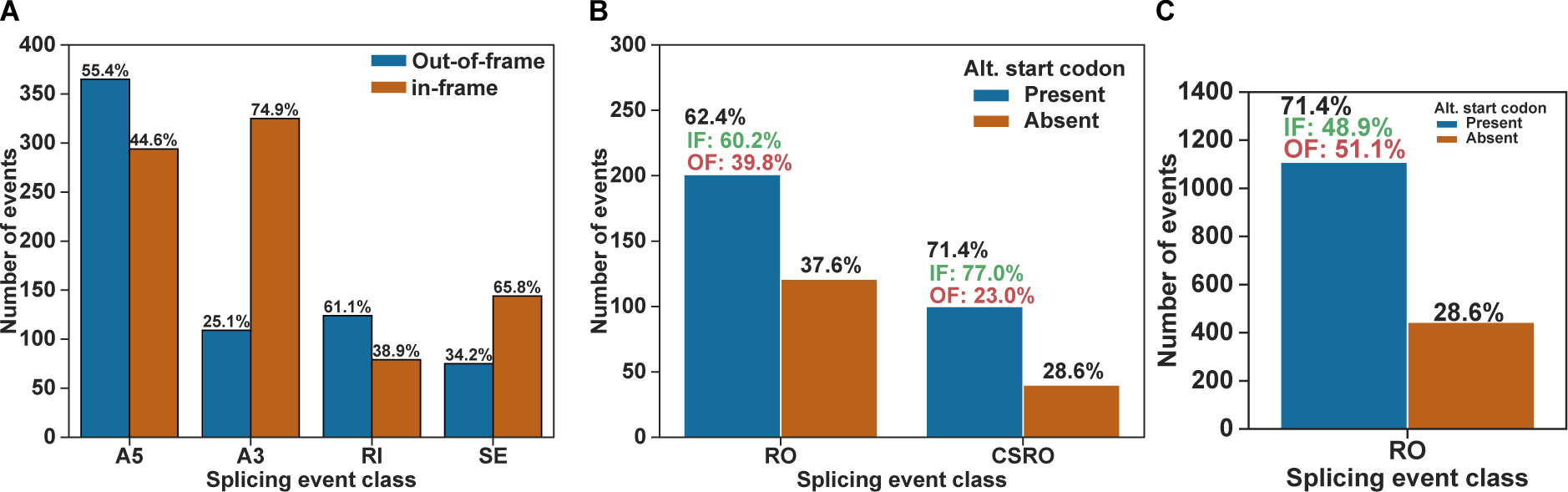
Alternative splicing events alter the protein coding potential of transcripts. **(A)** Bar plot showing the percentage of out-of-frame and in-frame changes by each class of cis-splicing events observed in *mett-10* mutant animals. **(B)** Bar plot showing the presence or absence of alternative start codon within the outron retained regions observed in *mett-10* mutant animals. The percentage of in-frame and out-of-frame start codons are shown above the blue bars. **(C)** Presence or absence of alternative start codons within a 120nt window upstream of annotated canonical start codons across *C. elegans* genes. The percentage of in-frame and out-of-frame start codons is shown above the blue bar.

For trans-splicing events, we specifically looked for potential upstream open reading frames that can arise due to outron retention or the cis-splicing of retained outrons. For outron retained and cis-spliced retained outron events, we used the most extended Nanopore Direct RNA sequencing reads as representatives of the 5′ end of transcripts. In most cases, these reads will likely be shorter than the actual transcription start sites (42). Next, we searched for the ATG start codon within outron regions or the cis-spliced retained outron regions. We found that 62.4% of all retained outron regions and 71.4% of all cis-spliced retained outrons contain at least one start codon sequence (Figure 8B). In both outron retention classes, most potential upstream start codons are in-frame with the downstream canonical start codon (Figure 8B). The percentages of in-frame start codons within outron regions of outron-retaining genes are higher than in-frame start codons observed across all *C. elegans* genes when considering a median outron size of 120nt (Figure 8C). Therefore, *mett-10* sensitive outron regions are more likely to contain an in-frame upstream start codon, and outron-retained transcripts could be translated into novel protein isoforms.

## Discussion

### Absence of U6 snRNA m6A43 in *mett-10^-/-^* animals leads to a wide range of cis-splicing defects

The cellular and molecular function of many snRNA modifications remains elusive. U6 snRNA m6A modification at A43 and the U6 snRNA sequence where the modification is found are highly conserved from *S. pombe* to humans. The U6 snRNA sequence UACAGA is essential for both the methylation reaction by METT-10 / METTL16 and the 5′SS recognition during pre-mRNA splicing (5, 68, 70). We identified that, in *C. elegans,* the absence of *mett-10* causes large-scale cis- and trans-splicing defects compatible with the function of METT-10 in m6A methylation of U6 snRNA. Previously, it was suggested that *mett-10* mutants do not show global splicing defects (23). We were able to capture the cis- and trans-splicing changes by deep sequencing of Illumina libraries in quadruplicate, using Nanopore Direct RNA sequencing and building condition-specific reference transcriptomes. Our results show that the pre-mRNAs of thousands of genes are mis-spliced in *mett-10* mutant animals, and many of these events are likely to contribute to the developmental and germline phenotypes of *mett-10* mutants.

We further show that, in *C. elegans,* U6 snRNA m6A43 functions to recognise 5′SSs with a //GURAG motif and when U6 snRNA is not m6A methylated, 5′SSs move to sites enriched with AG//GU (Figure 9A). In this context, the adenosine at the +4 position can distinguish 75% of all 5′SSs sensitive to the absence of U6 snRNA m6A43. Our results show that the mechanism of 5′SS recognition by U6 snRNAs that are m6A methylated at A43 is conserved between yeast, plants and animals (21, 24). U6 snRNAs with m6A43 potentially interact stronger with the 5′SSs with +4A through trans-Hoogsteen sugar edge interaction between m6A43:+4A (Figure S9G-I) (5, 21, 77). Without m6A43, 5′SSs move to sequences that support more robust U5 snRNA recognition, such as AG//GU, a stronger unmethylated U6 snRNA recognition, such as +4U, or both (Further discussed in (21, 24, 75)). Consequently, it is possible that spliceosome aborts splicing altogether where a suitable 5′SS is not found, as in intron retention and exon-skipping. Indeed, the absence of U6 snRNA m6A43, for many transcripts, leads to intron retention and exon-skipping where 5′SS //GURAG motif and +4A are essential determining factors. Importantly, there is no correlation between splicing changes and gene expression (Figure S9A and S9B), further supporting a direct role for *mett-10* and U6 snRNA m6A43 in splicing regulation.

**Figure 9.**
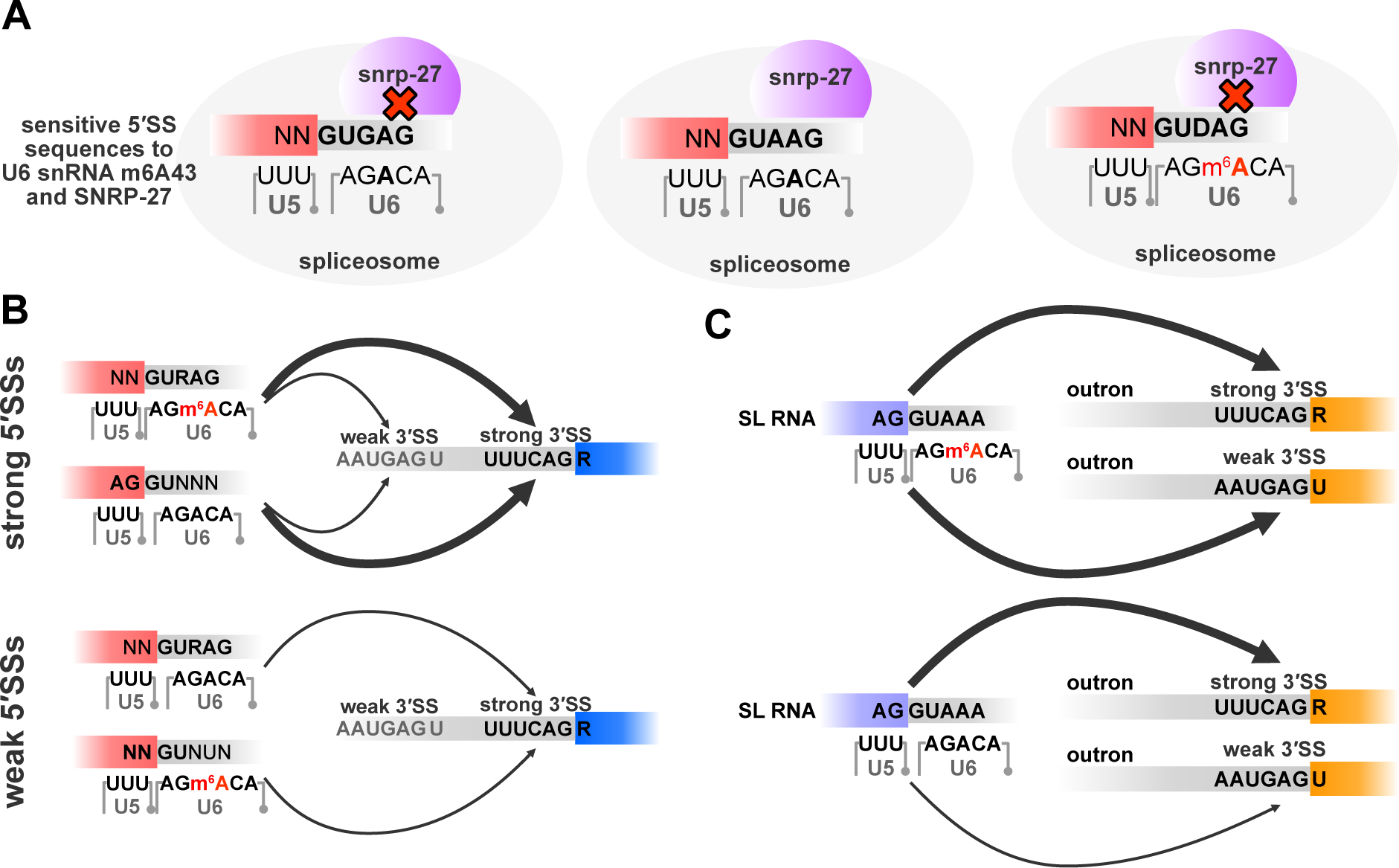
Model for splice site regulation by U6 snRNA m6A43, SNRP-27 and the splice site sequences. **(A)** U6 snRNA m6A43 and SNRP-27 function together to effectively and accurately recognise 5′SSs with a +4A and +3 position shows specific preference for the presence and absence of U6 snRNA m6A43, SNRP-27 or both. **(B)** Strong 5′SS interactions at //GURAG sites maintained by U6 snRNA m6A43 or strong U5 interactions can support the usage of weak upstream 3′SSs (upper panel). When 5′SS interactions weaken, only strong 3′SS are used (bottom panel). **(C)** When U6 snRNA is m6A methylated, strong recognition of the 5′SS on the SL RNA can support the usage of weak 3′ trans-splice sites (upper panel). When U6 snRNA - 5′ trans-splice site interactions weaken, trans-splicing at weak 3′ trans-splice sites fail (bottom panel).

### U6 snRNA m6A43 and SNRNP27K functionally interact for accurate 5′SS recognition but regulate distinct 3′SSs

Available structures of the spliceosome suggest that the spliceosomal protein SNRNP27K interacts with the U6 snRNA and could function in 5′SS recognition (Figure S9G) (5). Recent re-analysis of the cryo-EM structures of the human spliceosome suggests that M141 of SNRNP27K is positioned close to the m6A43 of U6 snRNA (75). Our analysis of the available RNA-Seq data from SNRP-27 M141T mutants strongly supports a functional interaction between U6 snRNA m6A43 and SNRP27K in recognition of 5′SSs with +4A (Figure 9A). The effect of U6 snRNA m6A43 and SNRP-27 on alternative usage of 3′SSs is distinct. In the absence of U6 snRNA m6A43, 3′SS usage shifts from weak upstream 3′SSs to downstream strong canonical 3′SSs, suggesting that U6 snRNA m6A43 is important for the stabilisation of 5′ and 3′SS interactions (Figure 9B). Our data suggests that, in *C. elegans*, this interaction could depend on the strength of 3′SS sequences primarily based on U2AF binding motifs. In the absence of SNRP-27, usage of strong downstream 3′SSs shifts to weaker upstream positions, suggesting that in the absence of SNRP-27, distance of the 3′SS to the 5′SS could be a critical factor and SNRP-27 could function to facilitate distal 3′SS selection. U6 snRNA interacts with both the 5′SS and 3′SS during the C to C* transition of the spliceosome (78), as was discussed for the *A. thaliana* alternative 3′SS usage (21), and this interaction can explain why U6 snRNA m6A43 is important for 3′SS selection. Indeed, many spliceosomal proteins associated with the C* complex affect 3′SS usage (79). However, SNRNP27K is no longer present in human C and C* complexes. It is possible that the impact of SNRP-27 on orienting the 5′SSs for U6 snRNA binding in *C. elegans* extends beyond the time it is present in the spliceosome.

In *C. elegans*, similar shifts in alternative 3′SS usage events have been observed in tissue-specific gene expression (80) and during ageing (81). Therefore, U6 snRNA m6A43 and SNRP-27 could have regulatory roles during animal development.

### The diversity of 5′SS sequences suggests U6 snRNA m6A methylation could be a regulated process

Overall, our analysis of splicing events observed in *mett-10^-/-^* and wild-type animals suggests there are two classes of 5′SSs in *C. elegans*: 5′SSs with the //GURAG motif and +4A, which prefer U6 snRNAs that are m6A methylated and the 5′SSs with AG//GU or +4U that would prefer unmethylated U6 snRNAs. Our experimental data is consistent with the previous *in silico* analysis of annotated *C. elegans* 5′SSs (21). Indeed, within all *C. elegans* 5′SS sequences (Figure S9C), 44% of the 5′SSs have a //GURAG motif (Figure S9D), 32% of the 5′SSs have an AG//GU without //GURAG (Figure S9E), and 16% of the 5′SSs have a +4U (Figure S9F) which also strongly favour AG//GU. Our RNA-Seq analysis of *mett-10* sensitive 5′SSs and the presence of different 5′SS classes could mean not all U6 snRNAs are m6A methylated in cells, which could also be a regulated process or cells could use +4A and non-+4A 5′SSs to regulate the expression level of transcript isoforms.

### U6 snRNA m6A43 is required for SL trans-splicing

U2, U4, U5 and U6 snRNAs are all required for SL trans-splicing in nematodes. However, as the trans-spliceosome structure is unknown, we do not know how snRNAs interact with the 5′ and 3′SSs during SL trans-splicing. Nanopore Direct RNA sequencing allowed us to capture the 5′ ends of mRNAs at an isoform level, and we were able to detect and quantify SL trans-spliced and outron retaining full-length RNA reads. Our results show that U6 snRNA m6A43 is required for efficient SL trans-splicing, and most outron retention events are explained by weak 3′ trans-splice site sequences (Figure 9C). Therefore, our results suggest that U6 snRNA interacts with the SL RNA 5′SSs similar to pre-mRNA 5′SSs, and in the absence of U6 snRNA m6A43, interactions with the +4A at the SL RNA 5′SSs weaken. The importance of 3′ trans-splice site sequences in the absence of U6 snRNA m6A43 suggests that U6 snRNA interactions with both the 5′ and 3′SS are likely conserved in SL trans-splicing.

### The function of METT-10 in mRNA modification is unclear

*sams−3,−4 and −5* all have 5′SSs with weak U5 snRNA interacting sequences and a //GURAG motif, which could make these sequences sensitive to the loss of *mett-10*. This could shift the usage of weak upstream 3′SSs to the stronger downstream 3′SSs, as seen for most alternative 3′SS usage events. However, the alternative 3′SSs of *sams−3, −4 and −5* used with low frequency in wild-type animals have a strong UUUCAG/R motif (Figure S4). Therefore, METT-10 mediated m6A methylation at the pre-mRNA introns of *sams−3, −4,* and *−5* could be a unique event in *sams* genes.

There is biochemical data supporting *in vitro* m6A methylation of *sams* pre-mRNAs by METT-10 (23, 35, 82), but evidence for *in vivo sams* pre-mRNA m6A methylation is limited. Using the highly sensitive SCARLET method (83), Mendel et al. could detect significant U6 snRNA m6A methylation in total RNA, but m6A methylation was undetectable on *sams−3* pre-mRNAs (23). Watabe et al. used Oxford Nanopore Direct RNA sequencing with *in vitro* transcribed *sams−3 and −4* transcripts that are either m6A methylated or not and trained machine learning algorithms, which were then used to classify the endogenous *sams−3 and −4* transcripts in the non-sense mediated decay mutants (*smg-2)*. Their analysis suggests that most *sams−3/−4* transcripts are m6A methylated (35). However, this approach does not consider the transcriptome-wide signal differences in the Oxford Nanopore Direct RNA sequencing data. Our comparison of the Nanopore Direct RNA sequencing data between *mett-10^-/-^* and wild-type animals using two transcriptome-wide analysis tools did not reveal any significant sites of m6A modification within *sams* genes. Therefore, m6A modification of *C. elegans* mRNAs requires further investigation.

### The biological significance of the alternative splicing events

The majority of the splicing events, either cis- or trans-, have the potential to alter the open reading frame and, therefore, could significantly impact protein expression for hundreds of genes. Indeed, most intron retention and exon-skipping events generate out-of-frame changes. Although the majority of the alternative 5′SS events generate out-of-frame changes (55.4%), a significant portion remains in-frame (44.6%). On the other hand, most of the alternative 3′SS events are in-frame, supporting the observation that weak and less frequently used 3′SSs move to strong canonical positions in the absence of U6 snRNA m6A methylation.

Most of the retained outron sequences in *mett-10^-/-^*animals have alternative start codons. Although we do not know if the translation machinery utilises these upstream start codons, genes sensitive to outron retention are more likely to have in-frame start codons within the outron sequences. Therefore, outron retention could generate novel protein isoforms.

In summary, our results show that altering U6 snRNA function either through its m6A modification or through proteins that interact with U6 during spliceosome assembly, leads to multiple alternative splicing changes. While many of the alternative splicing events are likely to remove protein function, many other splicing changes reveal events that are regularly used in wild-type animals, further supporting a regulatory role for either the U6 snRNA m6A43 or the 5′SS +4 base composition.

## Data availability

All raw data related to RNA sequencing is deposited at the European Nucleotide Archive (https://www.ebi.ac.uk/ena/browser/home) under the accession number PRJEB65287.

## Acknowledgements

We would like to thank Dr Giulia Furlan and Dr Dhiru Bansal for their support during the early phases of the project, Dr David Wright for his help with some of the bioinformatics tools, the UEA School of Biological Sciences technicians and admin for their support in running our research laboratories, and Dr Rebecca Taylor for her comments on our manuscript. The research presented in this paper was carried out on the High Performance Computing Cluster supported by the Research and Specialist Computing Support service at the University of East Anglia. Some strains were provided by the CGC, which is funded by NIH Office of Research Infrastructure Programs (P40 OD010440). We thank Wormbase for providing access to the *C. elegans* genome, annotation and genetic resources.

## Author contribution

Aykut Shen, conceptualisation, data acquisition, data analysis, visualisation, investigation, review and editing; Katarzyna Hencel, conceptualisation, data acquisition, review and editing; Matthew T. Parker, conceptualisation, data analysis, visualisation, investigation of trans-splice site identification and SNRP-27 data analysis, advising for general data analysis; Robyn Scott, data acquisition, data analysis; Roberta Skukan, data acquisition, data analysis; Aduragbemi S. Adesina, data analysis; Eric A. Miska, conceptualisation, funding acquisition; Yunsun Nam, conceptualisation, data acquisition, data analysis, review and editing, funding acquisition; Wilfried Haerty, conceptualisation, funding acquisition, review and editing; Gordon G. Simpson, conceptualisation, data analysis, funding acquisition, review and editing; Alper Akay, conceptualisation, data acquisition, data analysis, visualisation, investigation, funding acquisition, original draft, review and editing.

---

**Figure S1.**
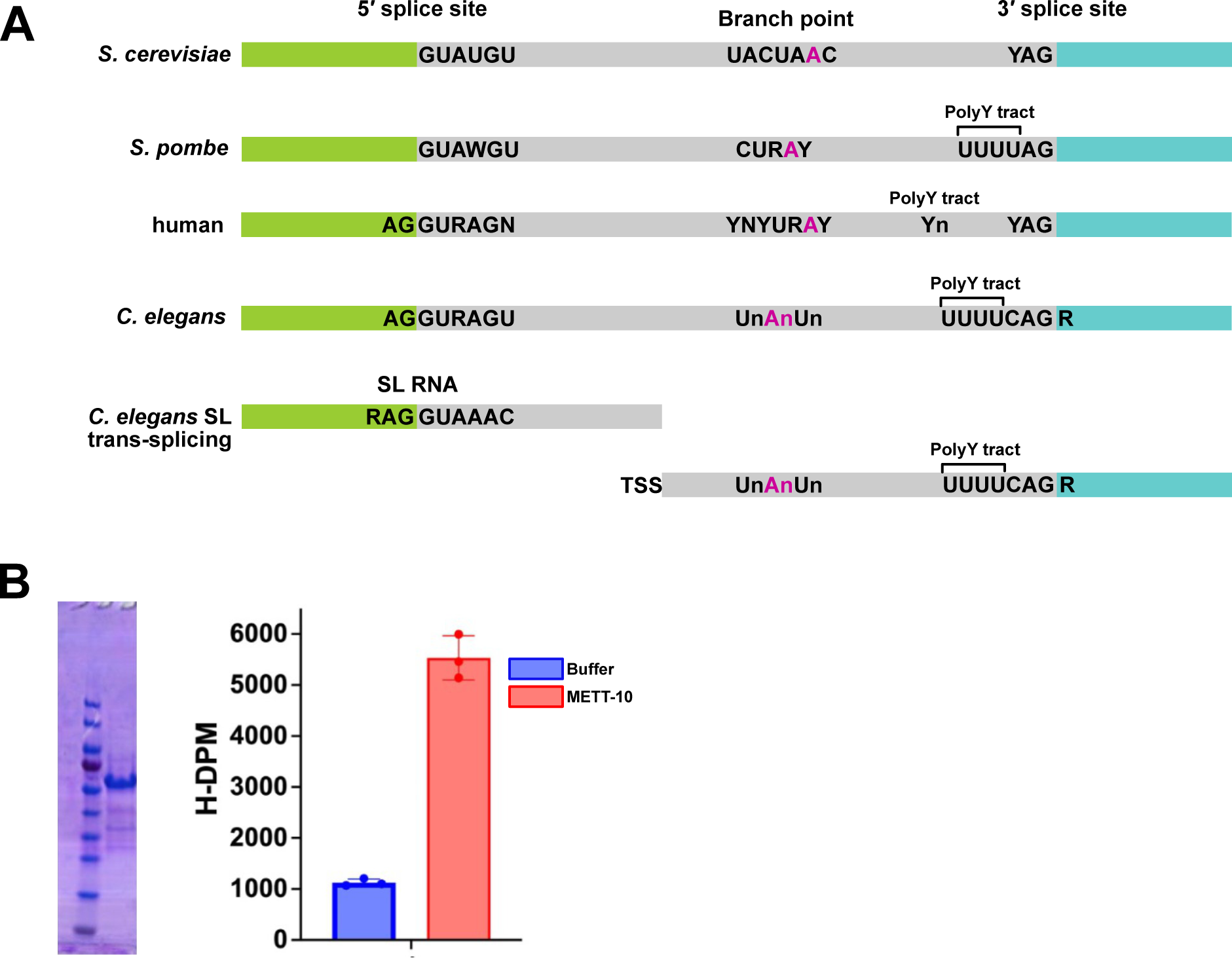
Splice site motifs in different organisms and methylation of U6 by METT-10. **(A)** 5′SS, 3′SS and branch point sequences in *S. cerevisiae, S. pombe, humans, C. elegans and C. elegans* trans-splice sites. 5′SS exons are shown in green, 3′SS exons in turquoise and introns in grey. Y: pyrimidines. TSS: transcription start site. **(B)** Gel electrophoresis of recombinant purified METT-10 (left panel), used in *in vitro* methylation reaction of U6 snRNA (right panel).

**Figure S2.**
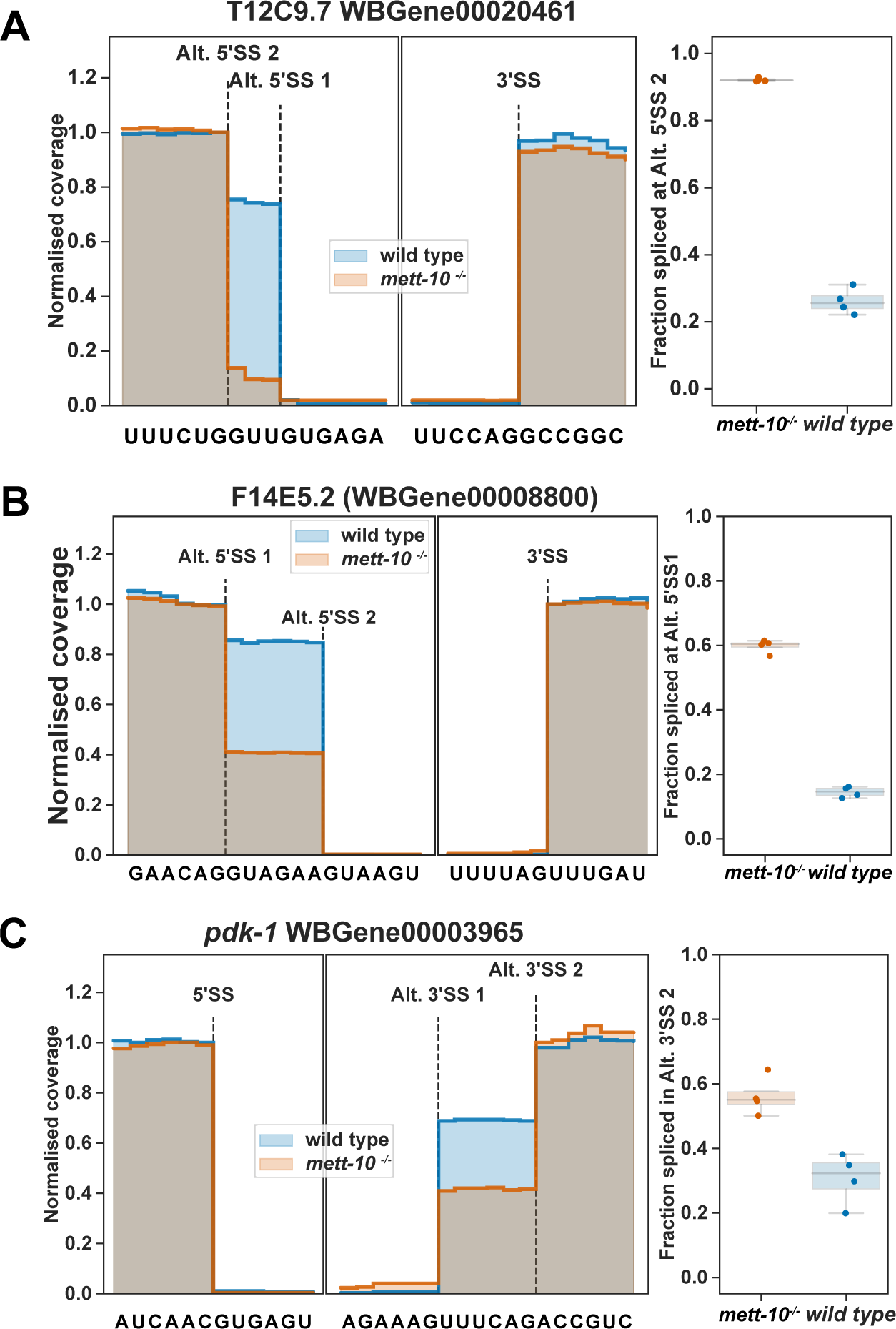
Additional examples of alternative 5′SS and 3′SS usage events. **(A)** T12C9.7 encodes the mitotic specific cyclin B2 and in wild type animals more frequently spliced at the Alt. 5′SS 1//GUGAG. In *mett-10^-/-^*, most of the splicing shifts to Alt. 5′SS 2 UG//GUUGU. **(B)** F14E5.2 encodes the *C. elegans* orthologue of the human GLG1 and F14E5.2 exon 8 is most frequently spliced at Alt. 5′SS 2 AA//GUAAG in wild-type animals. In *mett-10^-/-^* animals, 5′SS choice moves to the nearby Alt. 5′SS 1 AG//GUAGA. **(C)** *pdk-1* is the *C. elegans* orthologue of the human cancer associated kinase PDPK1. *Pdk-1* is frequently spliced in wild-type animals at both Alt. 3′SS 1 and Alt. 3′SS 2. In *mett-10^-/-^* animals most of the splicing events move to the Alt. 3′SS 2.

**Figure S3.**
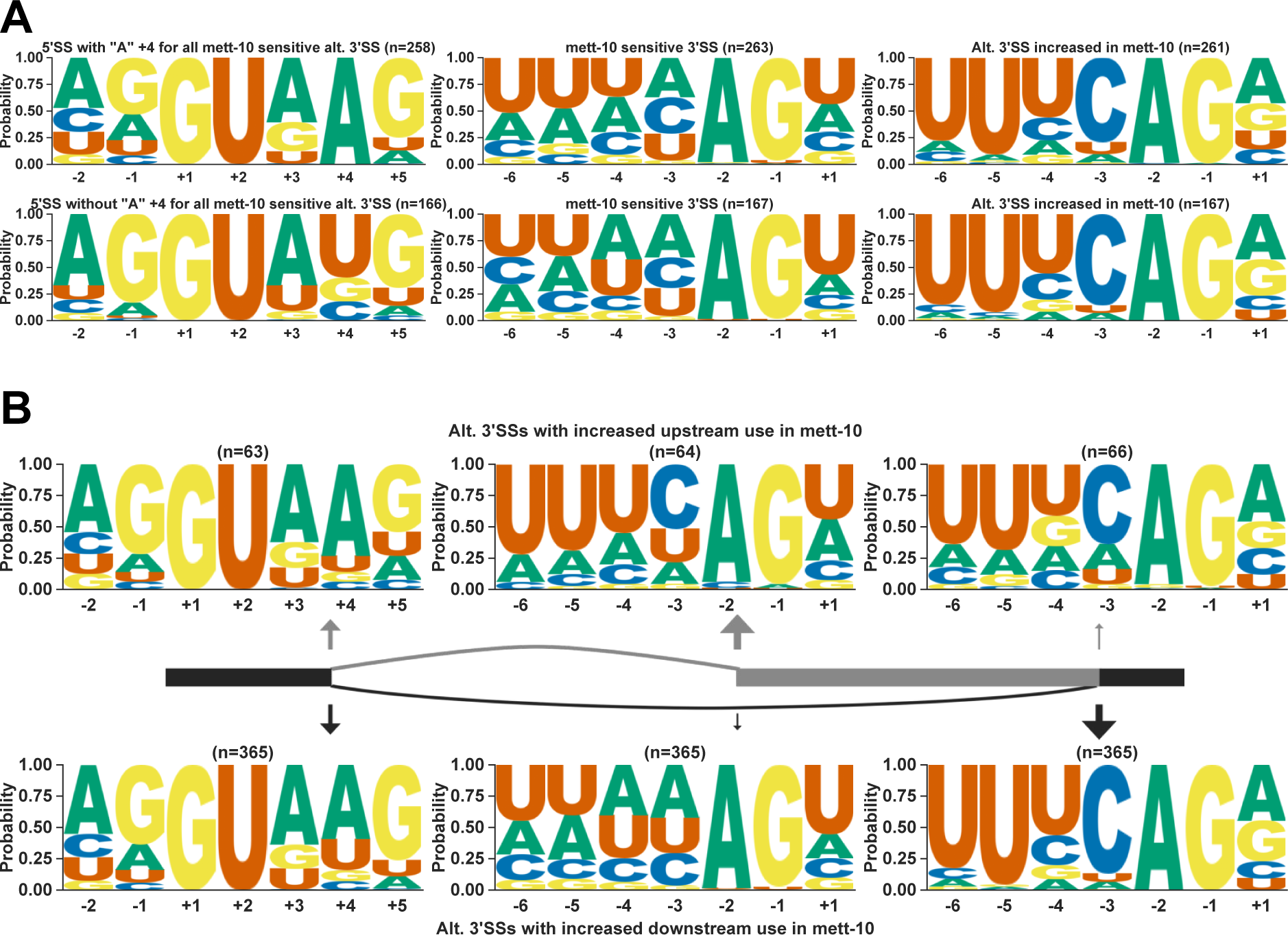
Alternative 3′SS usage. **(A)** Most alternative 3′SS events are associated with 5′SSs with +4A (263) as opposed to 5′SSs without +4A, which are enriched for AG//GU (167). *mett-10^-/-^* sensitive 3′SSs have weak 3′SS motif of UUUCAG/R and there is a switch from these 3′SSs to a strong 3′SS with a clear UUUCAG/R motif in *mett-10^-/-^*. **(B)** Majority of *mett-10^-/-^* sensitive 3′SS events (365) shift downstream to a stronger 3′SS motif (bottom panel). Alternative 3′SS events that shift upstream are limited (64) and tend to shift to a weaker 3′SS motif, although the difference is less clear.

**Figure S4.**
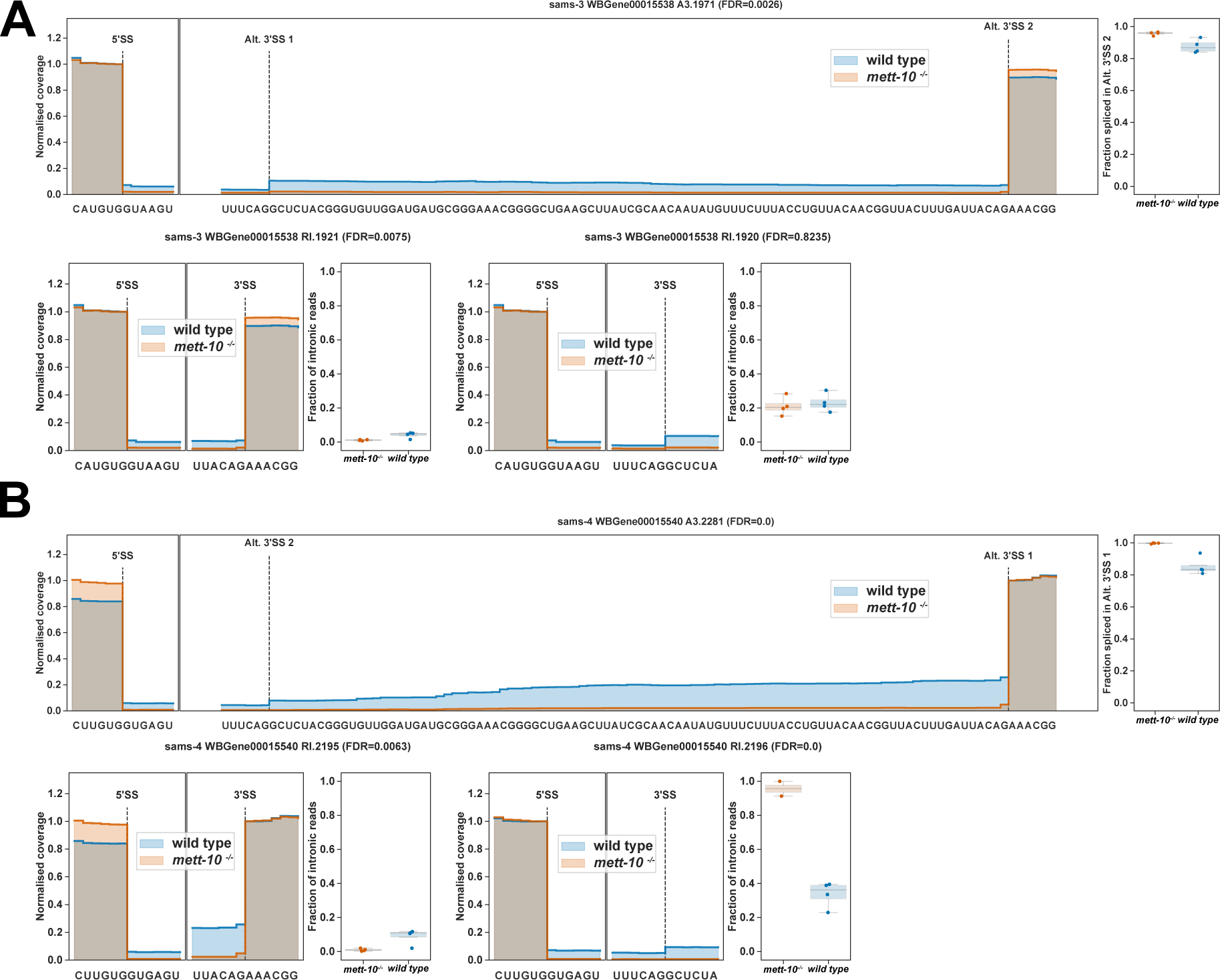
*sams* gene alternative splicing events. **(A)** RNA-Seq coverage of *sams−3* gene alternative splicing events for alternative 3′SS usage and two different intron retention events. **(B)** RNA-Seq coverage of *sams−4* gene alternative splicing events for alternative 3′SS usage and two different intron retention events.

**Figure S5.**
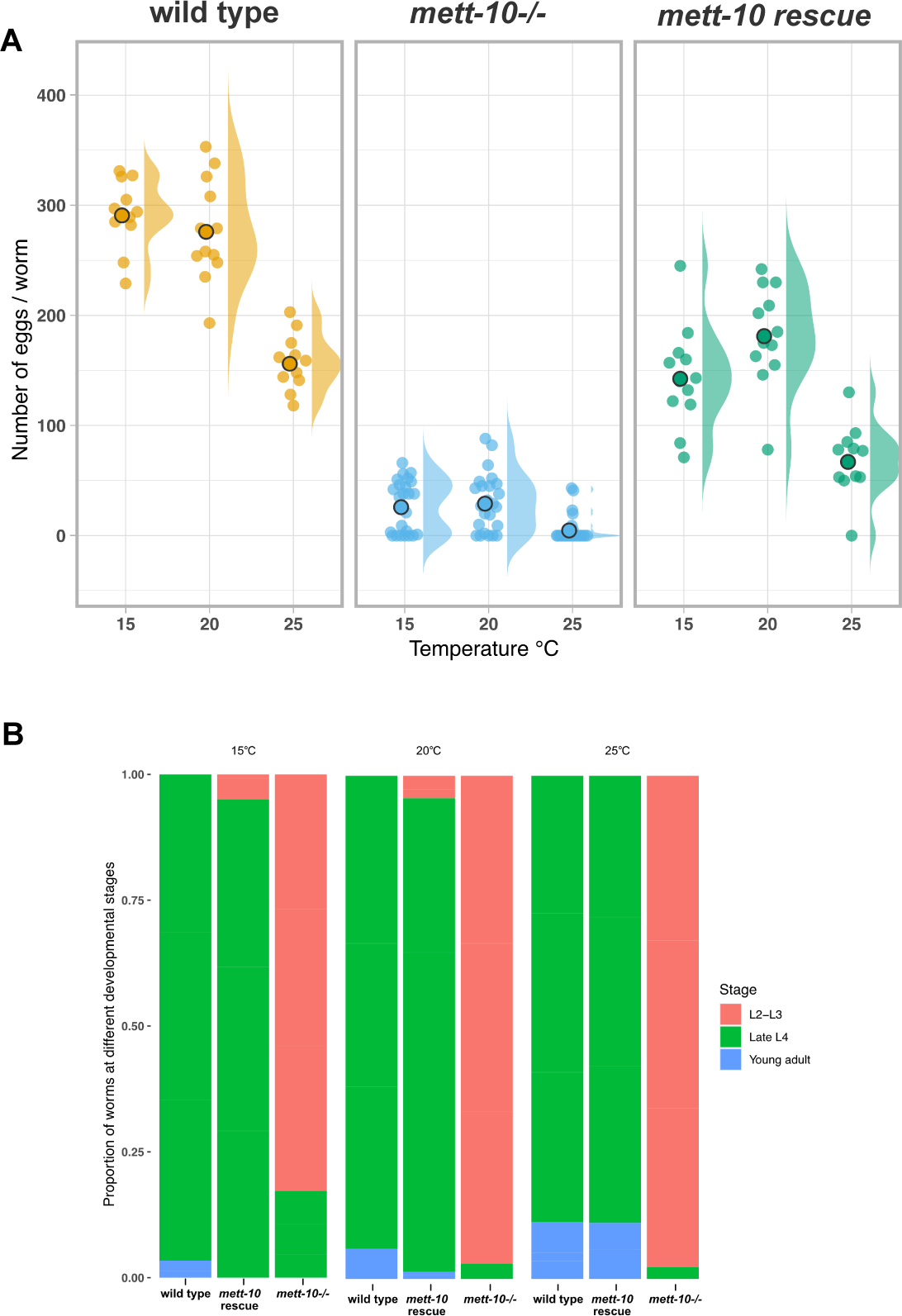
Phenotypic rescue of *mett-10(null)* animals with a germline expressed *mett-10* transgene. **(A)** Animals expressing the *mett-10 rescue* transgene in the germline and in a *mett-10-/-* background have higher progeny per animal at all 3 temperatures tested. **(B)** Animals expressing the *mett-10 rescue* transgene in the germline and in a *mett-10-/-* background have developmental timing similar to wild-type animals.

**Figure S6.**
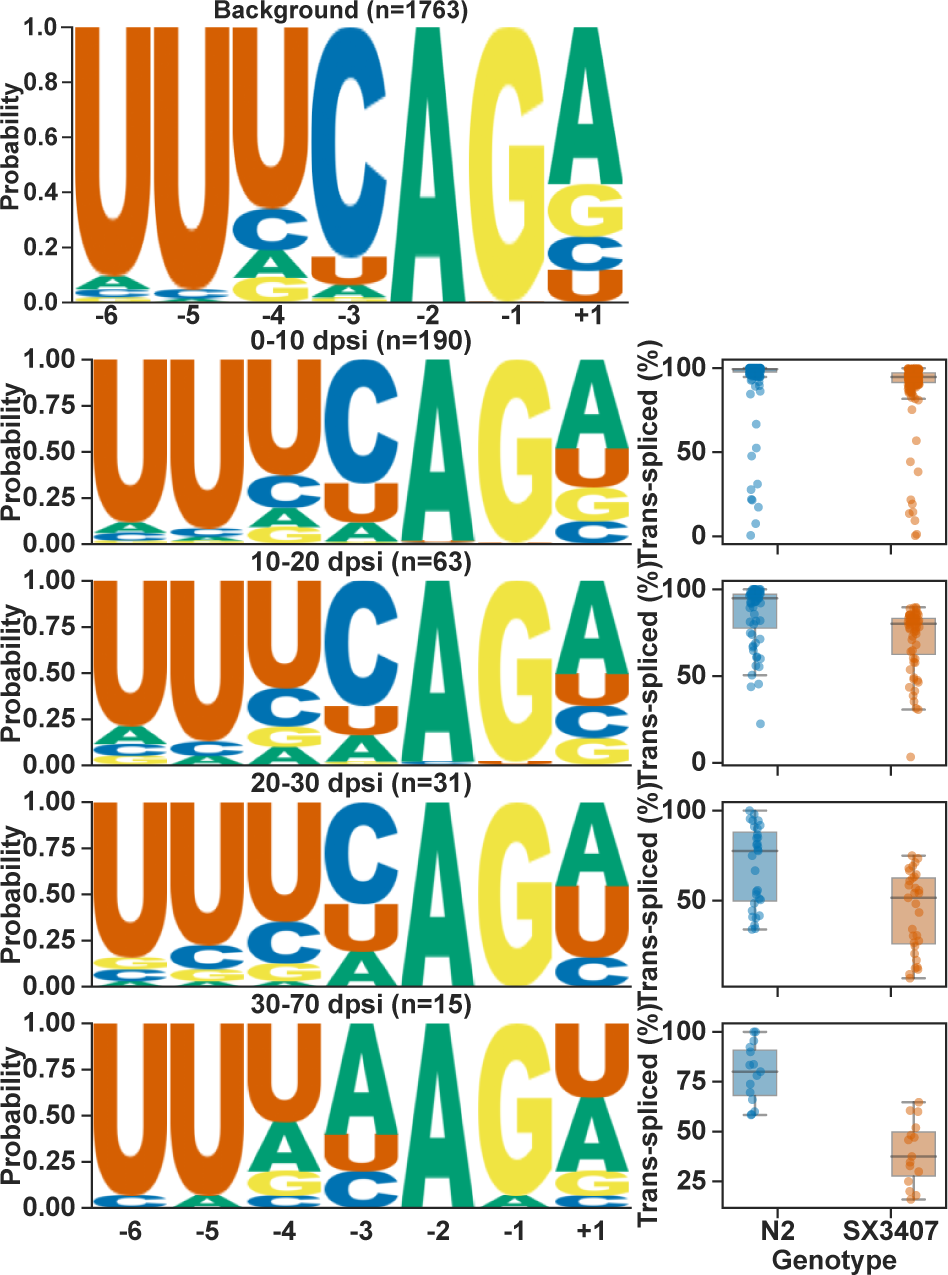
Analysis of 3′ trans-splice site motifs among transcripts with different levels of trans-splicing defects as in Figure 6A.

**Figure S7.**
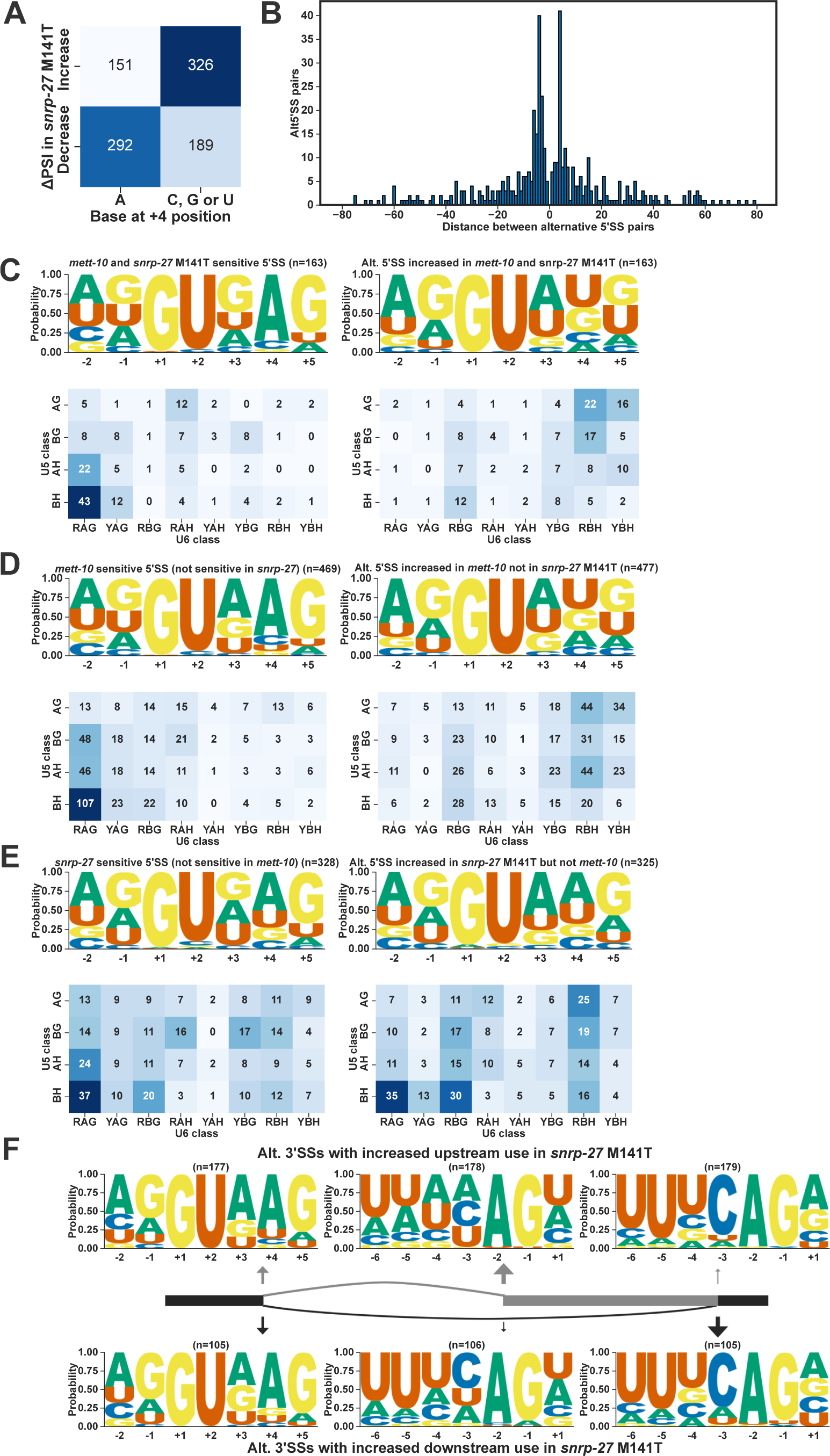
SNRP-27 is required for effective cis-splicing. **(A)** Heat-map showing presence or absence of +4A for *snrp-27* sensitive 5′SS events **(B)** Histogram of distance between alternative 5′SS pairs and the number of events. **(C, D and E)** 5′SS sequence motif and the frequency U5 and U6 interacting sequences for 5′SSs sensitive to **(C)** both *mett-10* and *snrp-27*, **(D)** only to *mett-10* and **(E)** only to *snrp-27.* **(F)** Sequence motif 3′SSs that are sensitive to *snrp-27* and the alternative 3′SS usage either shifts upstream (upper panel) or downstream (bottom panel).

**Figure S8.**
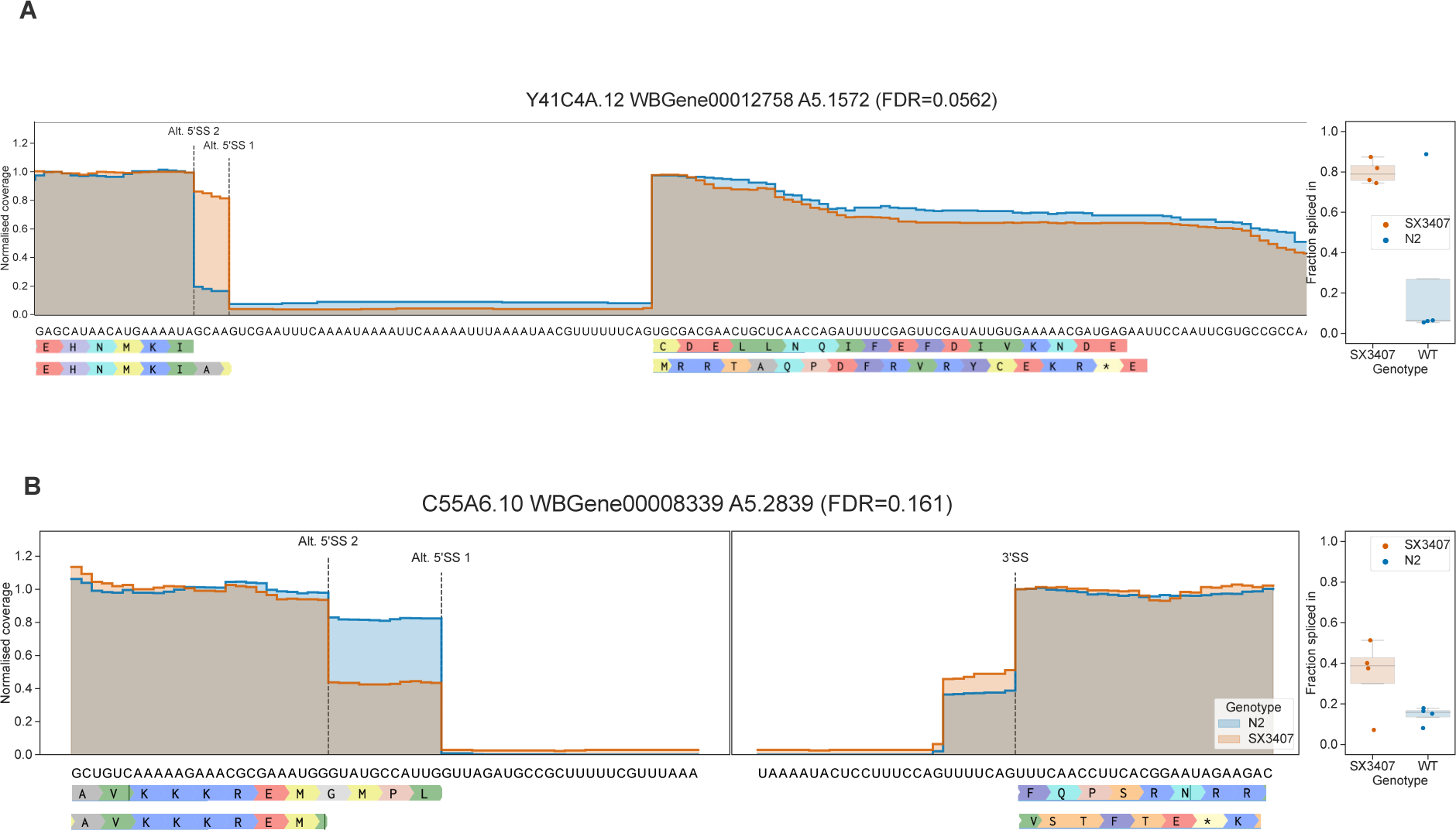
Examples of genes where alternative 5′SS usage generates a frame-shift in the open reading frame of **(A)** Y41C4A.12 and **(B)** C55A6.10.

**Figure S9.**
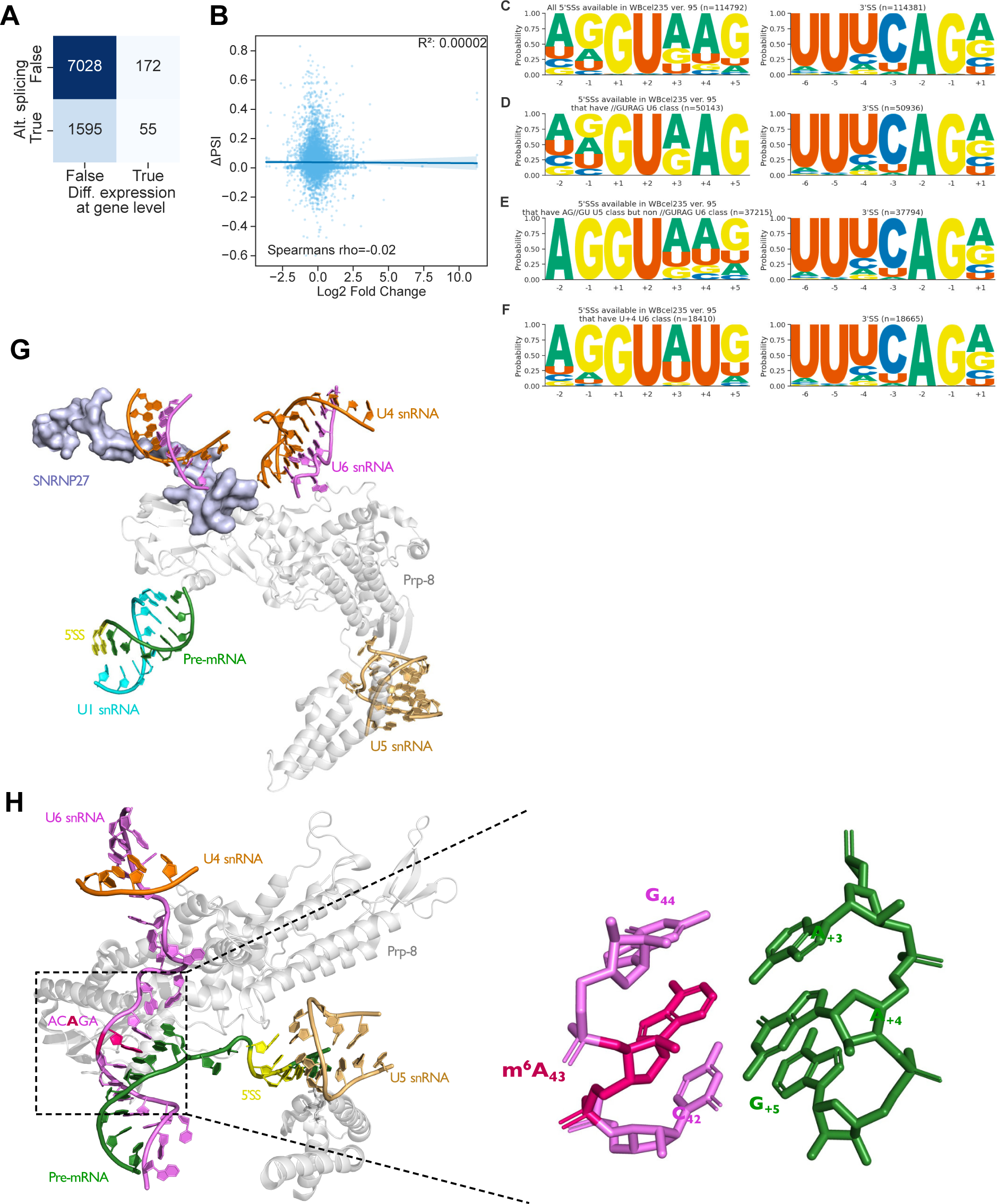
*mett-10* sensitive splicing changes do not correlate with the gene expression changes, sequence motifs of diverse splice sites in *C. elegans* and cryo-EM structures of pre-B and B complexes. **(A)** Heat map showing the overlap of genes with significant change in their expression and splicing. Only 55 genes with an FDR < 0.05 have a change in splicing and expression. **(B)** Correlation of splicing changes in *mett-10* mutants animals compared to wild type (ΔPSI) and the log2 fold change values. **(C-F)** Sequence motif of 5′SSs and 3′SSs for **(C)** all *C. elegans* genes, **(D)** with a //GURAG motif, **(E)** AG//GU and not //GURAG and **(F)** +4U only. **(G)** cryo-EM structure of the pre-B complex showing SNRNP27K, U6 snRNA, U1 snRNA, U5 snRNA and pre-mRNA PDB: 6QX9. **(H)** cryo-EM structure of the B-complex showing U6 snRNA - pre-mRNA 5′SS interactions PDB: 6AHD.

